# Organ-specific metabolic pathways distinguish prediabetes, type 2 diabetes and normal tissues

**DOI:** 10.1101/2021.05.09.443296

**Authors:** Klev Diamanti, Marco Cavalli, Maria J. Pereira, Gang Pan, Casimiro Castillejo-Lopez, Chanchal Kumar, Filip Mundt, Jan Komorowski, Atul Shahaji Deshmukh, Matthias Mann, Olle Korsgren, Jan W. Eriksson, Claes Wadelius

## Abstract

Defects in pancreatic islets and the progression of multi-tissue insulin resistance in combination with environmental factors are the main causes of type 2 diabetes (T2D). Mass spectrometry-based proteomics of five key-metabolic tissues on a cohort of 42 multi-organ donors provided deep coverage of the proteomes of pancreatic islets, visceral adipose tissue (VAT), liver, skeletal muscle and serum. Enrichment analysis of gene ontology (GO) terms built a tissue-specific map of the chronological order of altered biological processes across healthy controls (CTRL), pre-diabetes (PD) and T2D subjects. This unique dataset allowed us to explore alterations of entire biological pathways and individual proteins in multiple tissues. We confirmed the significant decrease of the citric acid cycle and the respiratory electron transport in VAT and muscle of T2D and we provided a thorough visual representation of the complete set of downregulated proteins. Importantly, we found widespread novel alterations in relevant biological pathways including the increase in hemostasis in pancreatic islets of PD, the increase in the complement cascade in liver and pancreatic islets of PD and the elevation in cholesterol biosynthesis in liver of T2D. Overall, our findings suggest inflammatory, immune and vascular impairments in pancreatic islets as potentially causal factors of insufficient insulin production and increased glucagon levels in the early stages of T2D. In contrast alterations in lipid metabolism and mitochondrial function in the liver and VAT/muscle, respectively, became evident later in manifest T2D. This first multi-tissue proteomic map indicates the temporal order of tissue-specific metabolic dysregulation in T2D development.

## Introduction

Insufficient secretion of insulin from pancreatic β cells and poor sensitivity to insulin from multiple tissues are important for the development of type 2 diabetes (T2D) that, in turn, is often followed by late complications, disability, increased mortality and increased health care costs. High energy diet and limited physical activity leading to obesity, as well as, heritability are predisposing factors that lead to an increased risk of T2D (Chen et al., 2012; Prasad and Groop, 2015). Beside genetic factors, various cellular components such as proteins and metabolites have been reported to be consistently altered in T2D, partly driven by environmental factors (Bellou et al., 2018; Diamanti et al., 2019; Roden and Shulman, 2019). Several studies have associated the effects of insufficient insulin with specific cellular deregulations including glucose levels and lipid deposition in tissues, and fatty acid uptake and metabolism (Heilbronn et al., 2004; Kusminski et al., 2009; Nagle et al., 2009).

The major tissues for the development of T2D include pancreatic islets, visceral adipose tissue (VAT), skeletal muscle and liver, but they remain largely understudied (DeFronzo, 2009). Instead the majority of studies exploring T2D have focused on easily accessible tissues such as adipose tissue and biofluids that exhibit cellular and molecular alterations reflecting events that may take place in the other, primary tissues, thus providing indirect evidence (DeFronzo, 2009; Shin et al., 2014). Exploring alterations in biological pathways across the various metabolically relevant tissues and distinct states of T2D would provide an information-rich holistic view of the primary events leading to the disease.

Liquid chromatography (LC) coupled with mass spectrometry (MS) is a methodology that is routinely employed for the identification and quantification of proteins from tissue samples, thus ultimately aiming at discoveries of biological and medical relevance (Aebersold and Mann, 2016). Technological advances including sample preparation, data acquisition and computational processing pipelines have enhanced the overall analytical capacity of MS proteomics (Bruderer et al., 2017; Kelstrup et al., 2018). MS proteomics has been utilized in a spectrum of tissues to study multiple diseases spanning from cerebrospinal fluid for Alzheimer’s disease, to brain for medulloblastoma and induced pluripotent stem cells differentiated into myoblasts for T2D (Archer et al., 2018; Bader et al., 2020; Batista et al., 2020).

In this study, we used tissue samples from 43 multi-organ donors and MS-based proteomics to build a map of altered biological processes in pre-diabetes (PD) and T2D in VAT, liver, skeletal muscle, pancreatic islets and serum. This resource provides an extensive coverage of the proteome of multiple tissues relevant to T2D. We confirmed a substantial fraction of well-characterized markers from the literature such as the significant downregulation of the citric acid (TCA) cycle in VAT and skeletal muscle, but more importantly, we identified various novel biological signals and pathways through comparisons across tissues that substantially expand the knowledge on early and overt T2D. A network of comparisons among healthy controls (CTRL), PD and T2D provided a chronological overview of responses of tissues and highlighted tissue-specific patterns of altered biological processes. In summary, we provide a unique resource of protein levels and enriched biological processes in the most important tissues involved in the development of T2D.

## Results

We performed a large proteomics analysis of five key-metabolic tissues including visceral adipose tissue (VAT), liver, skeletal muscle, pancreatic islets and serum, across 43 multi-organ donors (Figure 1). Organ donors were characterized by normoglycemia (n=17), pre-diabetes (n=14) and type 2 diabetes (n=12). Tissue-samples were obtained from The Nordic network for Clinical islet Transplantation, supported by the Swedish national strategic research initiative Excellence of Diabetes Research in Sweden (EXODIAB). Availability of all five tissues for the same donor was the primary selection criterion in a collection of more than 250 subjects (Figure 1a; Supplementary Tables S1 & S2). In total, we analyzed 195 samples and identified more than 20K proteins (Figure 1a; Supplementary Tables S2 & S3). Anthropometric characteristics including age, body mass index (BMI) and sex, showed minor associations to hemoglobin A_1c_ (HbA_1c_), glucose stimulated insulin secretion (GSIS) in pancreatic islets and T2D (Supplementary Table S1), while a subset of them was associated with the variance (median explained variance ≥1%) of a fraction of proteins (Supplementary Figure S1; Supplementary Table S4). Additionally, cold ischemic time (CIT) that has been reported to affect the phosphoproteome and potentially the proteome, as well as the days of hospitalization in the intensive care unit (ICU) were associated to the variance of proteins, while, surprisingly, this did not apply to the medications that were administered during the ICU hospitalization (Supplementary Figure S1; Supplementary Table S4) (Mertins et al., 2014). As expected, HbA_1c_ and GSIS were significantly associated with PD and T2D (Supplementary Table S1), and dimensionality reduction on protein abundancies showed limited subgrouping of the subjects (Supplementary Figure S2).

**Figure 1:**
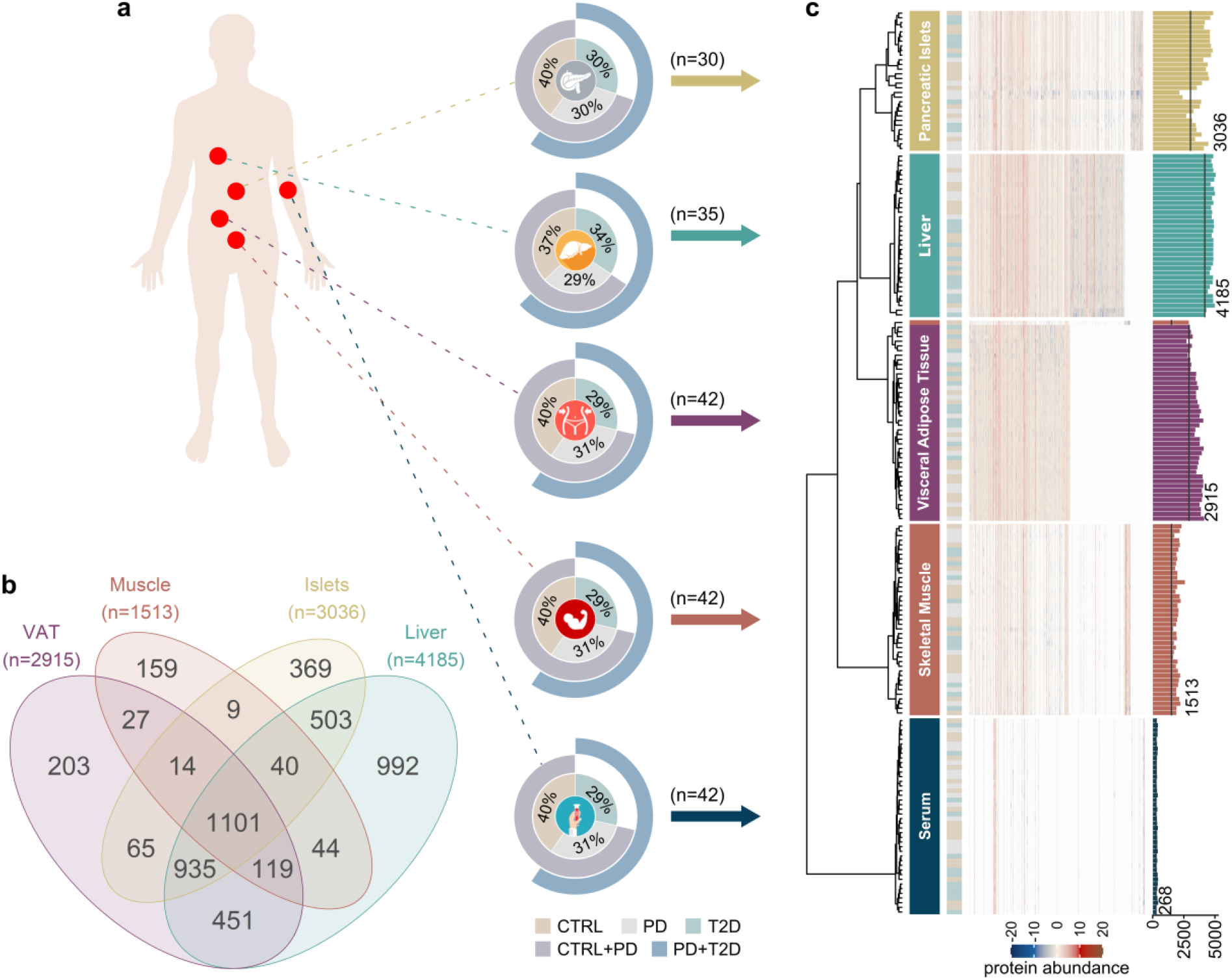
Overview of the study and exploration of the proteomics dataset. **a)** Schematic overview of tissue samples. Doughnut charts show the distribution of CTRL, PD and T2D subjects. The two outer circles show the fraction of subjects belonging to the merged groups of CTRL+PD and PD+T2D. **b)** Venn diagram that summarizes tissue-shared and tissue-specific proteins. A similar diagram including serum is shown in (Supplementary Figure S6a). **c)** Hierarchical clustering of protein abundancies across samples. Proteins were clustered using the function *ward*.*D* from the R package *stats* based on *1-r*, where *r* is the Pearson correlation coefficient. The dendrogram and the heatmap illustrate clustering and protein intensity, respectively. Tissues are indicated by color and text on the right-hand side of the dendrogram and the T2D status is shown in the next column. The right-most barplot shows number of identified proteins in each sample prior to filtering and the black line marks the number of proteins retained after filtering.

### MS proteomics allows accurate clustering of tissues

We performed a stringent quality control of samples and proteins to identify and remove potential outliers and sources of bias. Samples from the PD subject p14 were excluded due to consistently deviating more than two times from the median expression of proteins of the same tissue, in at least three tissues (Supplementary Figure S3) and the dataset was further processed for downstream analysis (Figure 1a; Supplementary Figures S4 & S5). We applied a conservative quality control approach for proteins to be identified in at least 80% of the samples of the same tissue, that resulted in removing more than 40% of the identified proteins across tissues; liver retained more than 4,000 of the originally identified proteins, pancreatic islets and VAT circa 3,000, and skeletal muscle more than 1,500 (Figure 1b). As expected, due to the vast dynamic range of the expressed proteome, serum had the lowest number of identified proteins, mainly, due to high abundance of some proteins (Geyer et al., 2017) (Figure 1b; Supplementary Table S2).

MS-based proteomics resulted in an extremely deep coverage of the proteome in tissue samples overcoming known challenges in VAT and skeletal muscle. We also achieved an impressive representation of the proteome of pancreatic islets that to date lacks extensive investigation (Brackeva et al., 2015; Metz et al., 2006). Proteins identified in single tissues (Figure 1b; Supplementary Figure S6a) showed strong enrichment for biological pathways and gene ontology (GO) biological processes highly relevant to the corresponding tissues. Proteins identified only in liver showed enrichment for metabolism of various types of lipids and amino acids (q<0.05), transport of bile acids and salts (q<6×10^−4^) and steroid biosynthesis (q<0.03) (Supplementary Figure S6b; Supplementary Table S5). Skeletal muscle was enriched for muscle-specific terms including muscle contraction (q<8×10^−3^) and development (q<0.03), as well as ion homeostasis (q<7×10^−3^), while pancreatic islets were primarily enriched for secretion of hormones (q<0.01) and insulin (q<0.04), and release of signals (q<3×10^−3^), and serum was enriched for the complement and coagulation cascades (q<0.05) (Supplementary Figure S6b; Supplementary Table S5).

Overall, a set of approximately 1,100 proteins was identified in all tissues excluding serum (Figure 1b), resulting in a collection of over 600 enriched GO biological processes (q<0.05) (Supplementary Figure S7). The 87 proteins that were identified across all five tissues resulted in many enriched GO biological processes, despite other larger collections of proteins identified among other subsets of tissues (Supplementary Figures S6a & S7). Overall, as expected, proteins identified in two or more tissues represented more generic biological functions than the ones identified in single tissues, including post-translational protein modifications (q<10^−5^), exocytosis (q<0.01), generation of precursor metabolites and energy (q<6×10^−17^), tissue homeostasis (q<2×10^−5^) and regulation of cell cycle G2/M phase transition (q<9.6×10^−12^) (Supplementary Figure S7; Supplementary Table S6).

Unsupervised clustering revealed identical proteomics profiles for tissue samples from subjects of the current set that were analyzed in a pilot study following a different MS acquisition mode, confirming the robustness of the analysis and the high quality of the data (Supplementary Figure S8). Furthermore, hierarchical clustering for protein abundancies showed that the tissues of origin carried the strongest biological signature, over the clinical subgrouping of subjects into CTRL, PD and T2D (Figure 1c; Supplementary Figure S2). The most abundant proteins of the muscle sample p42 that clustered with VAT (Figure 1c; Supplementary Figure S9a) were related to coagulation and immune response, and were similar to the ones from a subset of VAT samples (Supplementary Figures S9b & S9c), while others that were more similar to other skeletal muscle samples expressed muscle-specific signals (Supplementary Figures S9b-S9d) collectively suggesting contamination of the sample.

### Landscape of altered biological processes in PD and T2D

We performed within-tissue differential analyses for pairwise combinations of CTRL, PD and T2D while accounting for BMI, age, CIT, days of hospitalization in the ICU and, when applicable, percentage of purity of the pancreatic islets sample (Supplementary Figure S1; Supplementary Table S7). Next, we performed a ranked-list enrichment analysis to explore the landscape of enriched GO terms across tissues and identify altered biological processes across T2D phenotypes (Figure 2a-2c; Supplementary Figures S10 & S11; Supplementary Table S8). When comparing PD and CTRL, the vast majority of enriched GO terms occurred in pancreatic islets, while the other tissues had a limited number of significantly altered biological processes, even though a considerable number was enriched in liver. The dominance of enriched biological processes in pancreatic islets was slightly reduced when T2D was compared to CTRL. At the same time, we observed multiple enriched GO terms in skeletal muscle and VAT (Figure 2d). Pancreatic islets showed a limited number of enriched terms when T2D was compared to PD (10.5%), while ∼70% of the same enrichment terms were covered by liver (Figure 2d). A similar fraction of enriched terms for skeletal muscle was observed in the T2D-CTRL and T2D-PD networks, while for liver and VAT the largest number of enriched processes was observed in the T2D-PD network, 70.3% and 15%, respectively. On the other hand, pancreatic islets were responsible for over 66% of the enriched GO terms in both PD-CTRL and T2D-CTRL (Figure 2d). Overall, the distribution of clusters of GO biological process across tissues varied for different combination of CTRL, PD and T2D (Supplementary Figure S11a). In contrast to VAT, muscle and pancreatic islets that demonstrated enriched GO terms covering similar biological processes, the liver projected an intermediate state with more diversely enriched groups (Supplementary Figure S11b). Across tissues, pairwise comparison of liver and pancreatic islets, as well as VAT and skeletal muscle demonstrated large similarities on the patterns of enriched GO terms (Supplementary Figure S11b).

**Figure 2:**
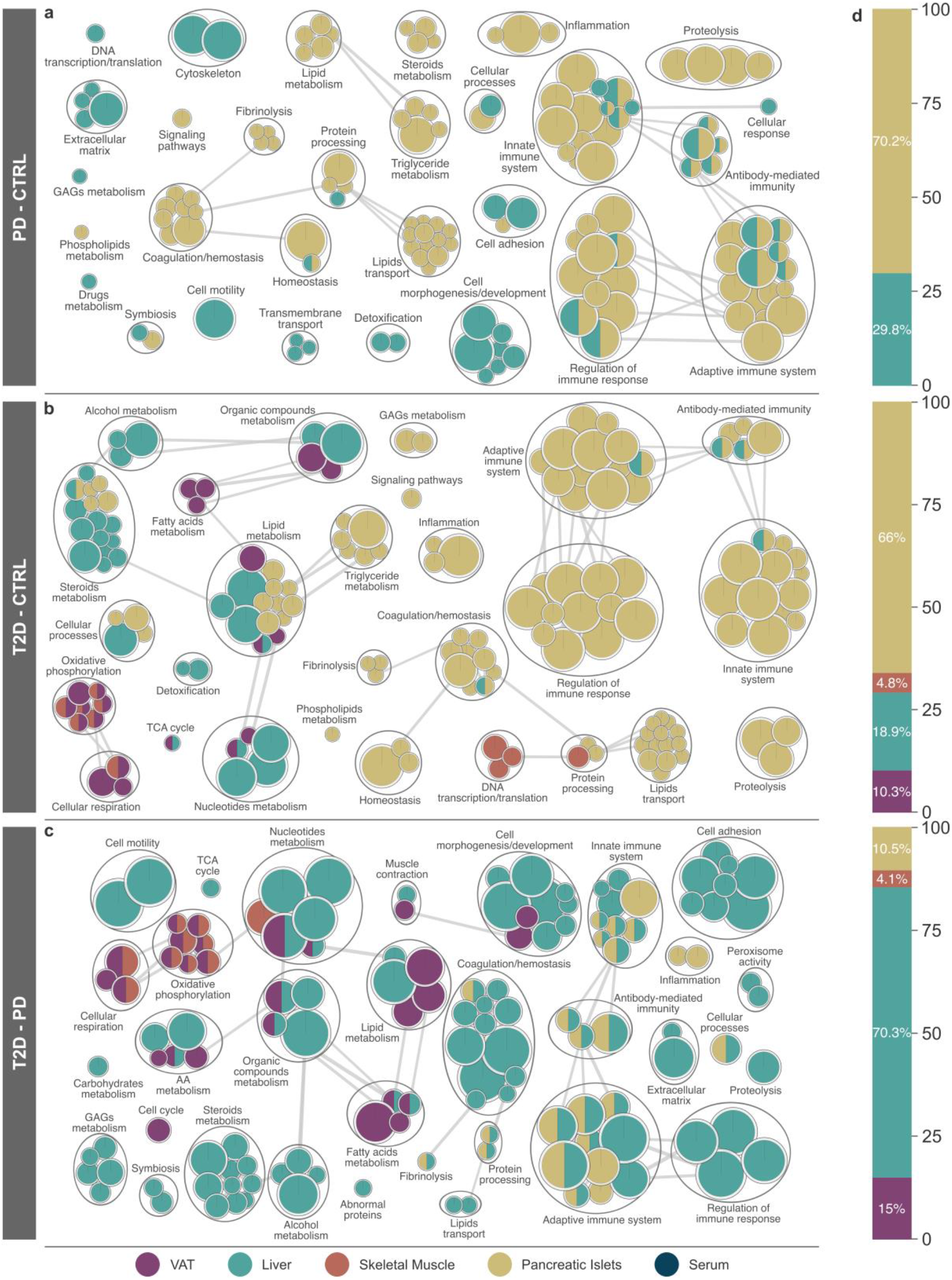
Overview of enriched GO biological processes across tissues for pairwise comparisons among CTRL, PD and T2D. The set of enriched terms was obtained as described in methods. **a-c)** Networks were created using the *Cytoscape* plugins *EnrichmentMap* and *AutoAnnotate* based on a curated set of custom classes for GO biological processes (Supplementary Table S8) (Merico et al., 2010). Nodes represent enriched GO biological processes (q<0.01) and their size indicates the number of proteins in the set. Thickness of edges represents the fraction of shared proteins between nodes and it was calculated using a Jaccard overlap combined index with *k*_*constant*_*=0*.*5*. Only edges describing high similarity (index≥0.7) were retained. Panels show comparisons of **a)** PD versus CTRL, **b)** T2D versus CTRL and **c)** T2D versus PD. **d)** Distribution of the enriched GO biological processes across tissues corresponding to the networks on the left-hand side. The distribution is presented as the fraction of enriched terms with respect to the total number of enriched terms for the pairwise comparison.

### Immune and lipid-related processes are upregulated in pancreatic islets of PD

The majority of enriched GO biological processes related to the immune system and its regulation in pancreatic islets was significantly upregulated in T2D (q_max_<3.7×10^−3^). This includes processes related to adaptive and innate immune systems, antibody-mediated immunity and inflammation were consistently elevated in pancreatic islets (q_max_ <7.1×10^−4^), as well as in liver (q_max_<5×10^−3^), of PD (Figure 2; Supplementary Figure S10). What is more, terms related to coagulation and homeostasis were also higher in T2D of pancreatic islets (q_max_<5.8×10^−3^) (Figure 2; Supplementary Figure S10). Homeostasis was significantly altered in PD of pancreatic islets (q_max_<2.5×10^−3^), both on the fibrinolytic and on the coagulation end, that when taken together with the elevated immune responses create the foundation of a pre-thrombotic state (Maschirow et al., 2015). Strong evidence of lipotoxicity including metabolism of steroids and triglycerides (q_max_<1.2×10^−3^), and metabolism of lipids as well as their transport (q_max_<2.5×10^−3^) were significantly increased in PD of pancreatic islets (Figure 2; Supplementary Figure S10).

### Alteration of biological processes in VAT and skeletal muscle of overt T2D

Biological processes connected to metabolism of steroids and alcohol were almost exclusively upregulated in liver (q_max_<8.4×10^−3^) whilst those associated with proteolysis increased in pancreatic islets (q_max_<4.7×10^−3^). We observed divergent directions in the alterations of the metabolism of organic compounds and nucleotides between liver (q_max_<6.7×10^−3^) and VAT (q_max_<4.8×10^−3^). We also identified a notable decrease in GO terms related to oxidative phosphorylation and cellular respiration in VAT (q_max_<4.1×10^−3^) and skeletal muscle (q_max_<3.4×10^−3^) (Figure 2; Supplementary Figure S10). The latter confirms our observation that enriched GO terms in VAT and muscle share a considerable fraction of semantics in T2D (Supplementary Figure S11b). Various processes related to the metabolism of fatty acids were exclusively decreased in VAT (q_max_<4.2×10^−3^), while skeletal muscle showed a significant decrease for terms related to DNA transcription and translation (q_max_<5.3×10^−3^) (Figure 2).

### Deregulation of various biological processes from PD to T2D in the liver

Comparison of T2D and PD resulted in the largest number of enriched biological processes across tissues with liver being responsible for over 70% of them and pancreatic islets for less than 11% (Figure 2). Immune-related terms and their regulatory processes largely occurred in liver (q_max_<6.3×10^−3^) that consists a shift in comparison to the former networks that compared PD and T2D to CTRL, and were dominated by pancreatic islets (Figure 2; Supplementary Figure S11a). This had twofold implications on the similarity of inflammatory and immune states between PD and T2D in pancreatic islets and on the late alterations of these biological processes in liver (Supplementary Figure S10). A large number of groups of enriched metabolic processes were exclusively altered in liver, probably due to its higher metabolic activity spanning multiple metabolic processes (q_max_<8.4×10^−3^). Specifically, metabolic processes of carbohydrates (q_max_<8.3×10^−3^), alcohol (q_max_<1.4×10^−4^), and steroids (q_max_<1.3×10^−3^) were upregulated, while metabolism of glycosaminoglycans (GAGs) was downregulated (q_max_<8.4×10^−3^) (Figure 2). Metabolic processes of amino acids (q_max_<8.2×10^−3^), organic compounds (q_max_<3.2×10^−4^), fatty acids (q_max_<3.3×10^−3^) and lipids (q_max_<7.7×10^−3^) were consistently upregulated in liver and downregulated in VAT (q_max_<2.2×10^−4^, q_max_<1.6×10^−7^, q_max_<2.1×10^−3^ and q_max_<2.8×10^−3^, respectively) of T2D in comparison to PD. Similarly to T2D-CTRL, oxidative phosphorylation and cellular respiration were enriched in muscle (q_max_<3.1×10^−4^ and q_max_<3.3×10^−5^, respectively) as well as VAT (q_max_<6.4×10^−4^ and q_max_<6.9×10^−5^, respectively), and followed the same direction of change (Supplementary Figure S10). Overall, biological processes enriched in the comparison T2D-PD resembled those enriched in T2D-CTRL in muscle and VAT. The latter did not apply to the same extent in liver, where about half of the biological processes were shared across comparisons (Supplementary Figure S11b).

### Enrichment of key metabolic pathways across tissues in PD and T2D

We analyzed the dataset for enriched biological pathways similarly to the aforementioned analysis for GO terms (Supplementary Table S9). We selected a subset of enriched biological pathways (q<0.05) that we grouped based on biological relevance (Figure 3). The TCA and its contributions of NADH and FADH_2_ electron carriers to the respiratory electron transport (RET), have been observed to be decreased in muscle of T2D and in adipose tissue of insulin resistant subjects (Befroy et al., 2007; Gaster, 2009; Heinonen et al., 2015). Here we observed that the entirety of TCA and RET were significantly decreased in T2D of VAT and muscle when compared to both CTRL and PD, as well as the merged group of CTRL+PD (Figures 3b & 3c; Supplementary Figure S12). At the same time, pathways related to the complement cascade were significantly increased in PD of liver and pancreatic islets (Figure 3a). In liver of T2D some of these pathways were significantly lower when compared to both CTRL and PD, suggesting widespread inflammation in PD and the inverse in T2D. Various resolution levels for biological pathways related to Fc-gamma and Fc-epsilon signaling, as well as hemostasis were significantly higher in PD and T2D in pancreatic islets, with the ones of T2D being significantly lower than the ones in PD. Metabolism of steroids enclosed five different pathways that were all significantly higher in T2D of liver as compared to CTRL, while differences between PD and CTRL did not show significance (Figure 3). Other enriched pathways included the increase of metabolism of vitamins in pancreatic islets of PD and T2D, and the decrease of pathways of neurodegenerative diseases in VAT and liver of T2D (Supplementary Figure S13). In the following sections we describe a set of representative enriched pathways in detail.

**Figure 3:**
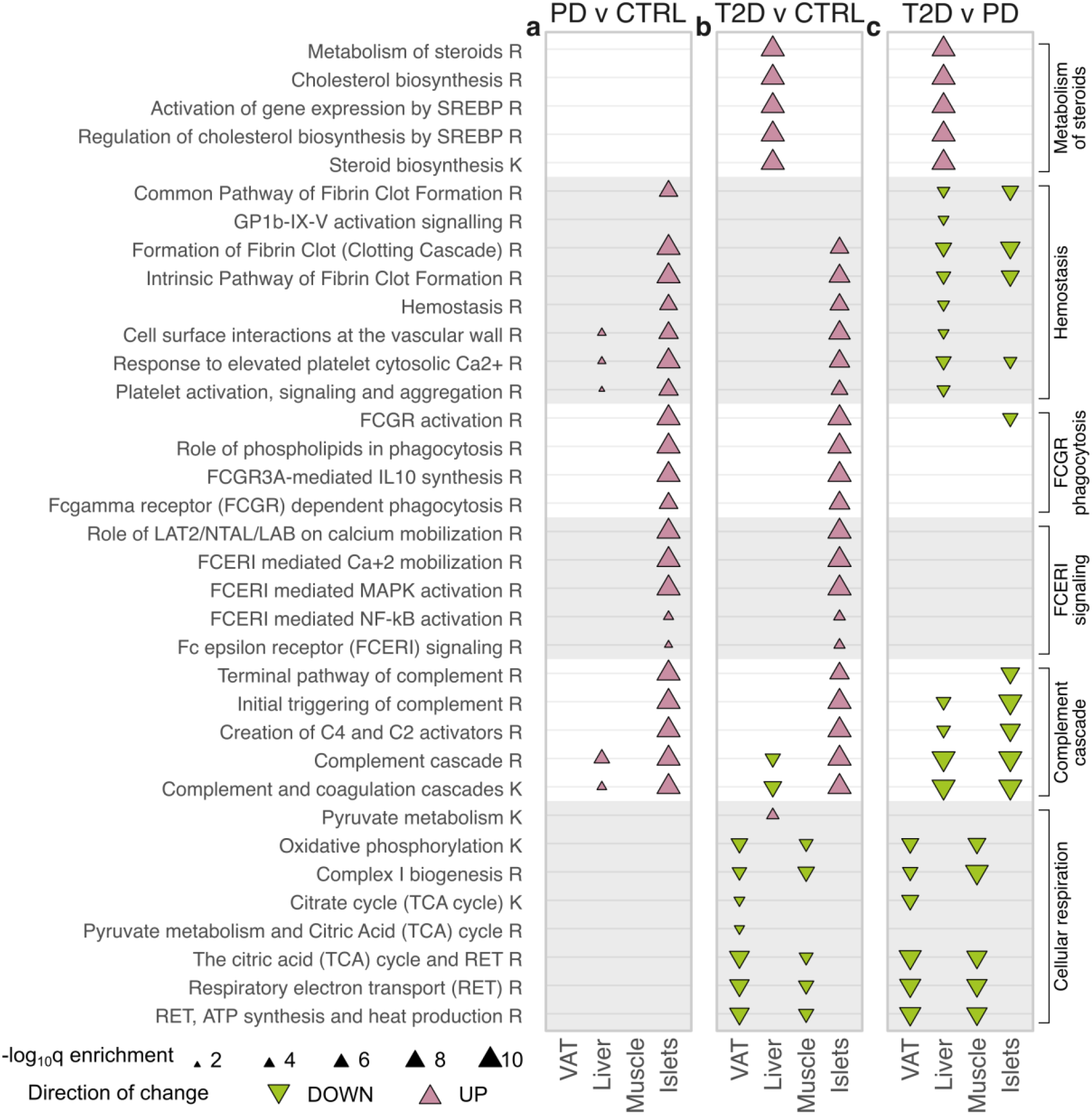
Significantly enriched biological pathways. Selected groups of enriched biological pathways (q<0.05) for pairwise comparisons among CTRL, PD and T2D. Specifically, **a)** PD-CTRL, **b)** T2D-CTRL, and **c)** T2D-PD. The set of enriched terms was obtained as described in the methods. Rows represent pathways, panels represent pairwise comparisons and columns within panels represent tissues. The database of origin for each pathway is noted with a suffix R or K for Reactome or KEGG, respectively. Sterol regulatory element binding proteins have been abbreviated as SREBP and Glycoprotein Ib-IX-V has been abbreviated as GP1b-IX-V.

### TCA and RET in VAT and skeletal muscle are significantly decreased in T2D

Mitochondrial function reflected by the TCA cycle and the RET chain were impaired in T2D in comparison to both CTRL and PD individuals (Figure 4). The same was also found for the biogenesis of the key mitochondrial component complex I, that is the leading component of the reduction in oxidative phosphorylation in skeletal muscle and VAT. In contrast, there were no evident differences in these pathways between PD and CTRL subjects. Analysis of mitochondrion-related GO cellular components revealed significant downregulation for VAT and skeletal muscle for several terms including mitochondrial matrix (q_VAT_<1.1×10^−5^ and q_muscle_<1.3×10^−2^), inner mitochondrial membrane protein complex (q_VAT_<7.2×10^−6^ and q_muscle_<1.7×10^−5^), mitochondrial envelope (q_VAT_<2.7×10^−6^ and q_muscle_<9×10^−5^) and mitochondrial protein complex (q_VAT_<10^−6^ and q_muscle_<1.7×10^−5^) (Supplementary Table S10).

**Figure 4:**
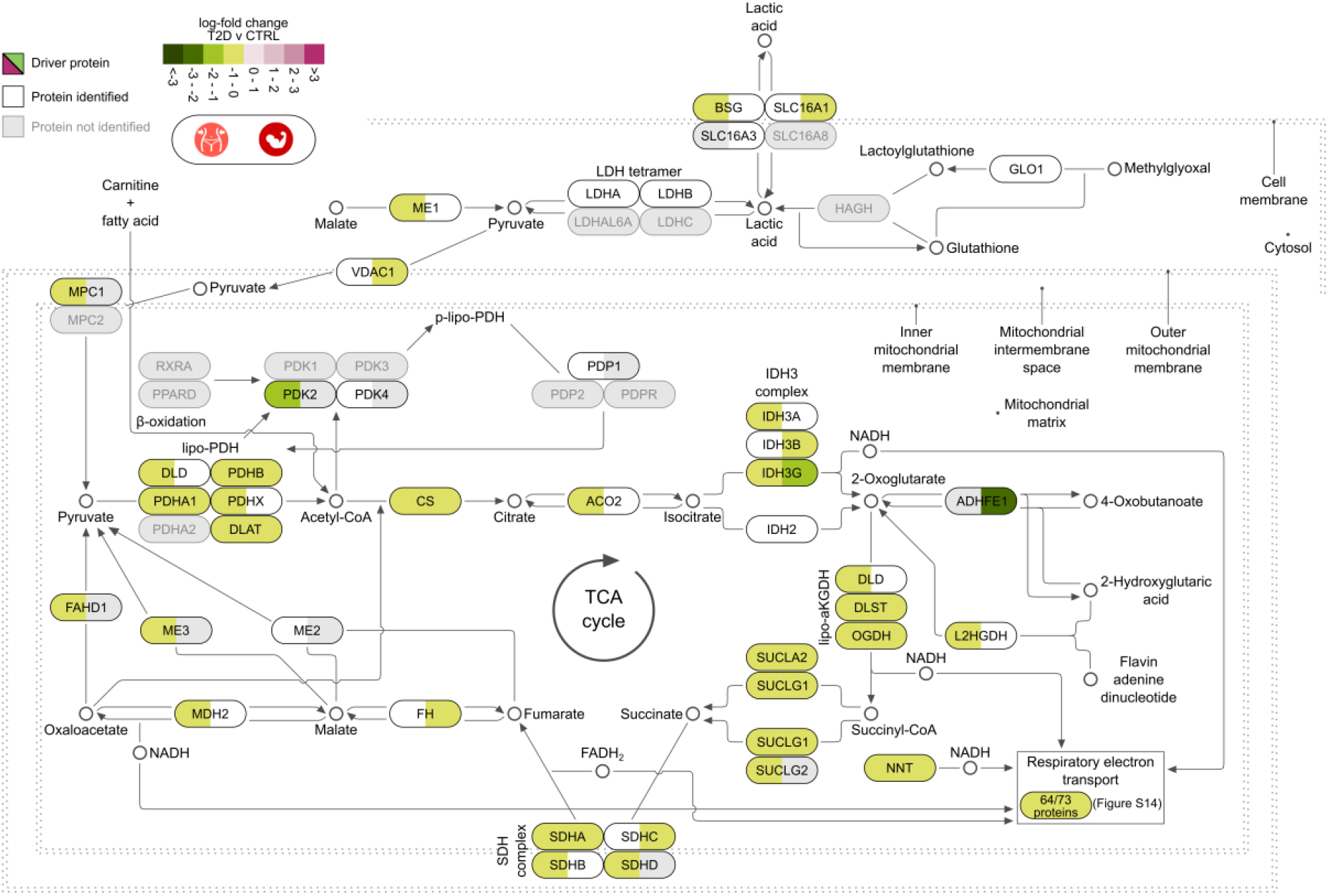
Biological pathway of pyruvate metabolism and the TCA cycle. The pathway is based on Reactome and complemented with details from KEGG and the literature. Tiles illustrate log-fold change values of T2D compared to CTRL in VAT (left hand side of tiles) and skeletal muscle (right hand side of tiles). Proteins leading the decrease of the pathway are marked as driving and are shown in black font and colored tiles. Proteins participating in the pathway and identified from MS proteomics, but not marked as leading edges are in white tiles, while those in grey tiles were not identified in our analysis.

The entirety of the TCA cycle and the RET chain was significantly lower in T2D than in CTRL in VAT and skeletal muscle (q<8.8×10^−8^ and q<3.2×10^−4^, respectively) (Figure 3; Supplementary Figure S8). The downregulation of the TCA cycle in VAT and muscle has been described in other studies to be associated with insulin resistance (IR) (Befroy et al., 2007; Gaster, 2009; Heinonen et al., 2015). The downregulation of pyruvate metabolism and the TCA cycle was driven by 26 proteins in VAT and 17 proteins in skeletal muscle, of which 11 were shared (Supplementary Figure S14). The vast majority of all 8 key enzymatic complexes of the TCA cycle, namely citrate synthase (CS), aconitase 2 (ACO2), isocitrate dehydrogenase (IDH), α-ketoglutarate dehydrogenase (αKGDH), succinyl-CoA synthase (SCS), succinate dehydrogenase (SDH), fumarate hydratase (FH) and malate dehydrogenase (MDH), were reduced in VAT and/or skeletal muscle of T2D, hence, confirming the significant downregulation of the entire pathway. The NADH and FADH_2_ electron carriers produced from the TCA cycle are subsequently imported to the RET and utilized for energy production (Figure 4). A large set of 54 proteins from VAT and 27 from skeletal muscle, of which 19 were shared, were marked to drive the overall decrease of the RET pathway components (Figure 3; Supplementary Figure S15). The majority of the driving proteins were moderately higher in PD than in CTRL, but their notable decrease in T2D resulted in the significant decrease of the pathway in both VAT and muscle. In conclusion, we confirm previous knowledge about the downregulation of TCA and RET in VAT and skeletal muscle in T2D. Furthermore, we extend this knowledge by identifying additional deregulated proteins, which highlights the value of the data.

### Cholesterol biosynthesis is significantly increased in liver of T2D

Multiple steps of the cholesterol biosynthesis pathway were enhanced in liver of T2D compared to PD as well as CTRL (Figures 3 and 5; Supplementary Figure S12b). The increase in liver of T2D subjects (q<1.6×10^−56^) was mainly driven by 16 proteins that demonstrated up to two-fold higher levels when compared to CTRL (Figure 5; Supplementary Table S9). Eleven of the 16 proteins are products of the biological pathway that activates gene expression through sterol regulatory element binding proteins (SREBPs) (Figures 3 and 5). Specifically, the expression of, among others, lanosterol demethylase (CYP51A1), 7-dehydrocholesterol reductase (DHCR7), farnesyldiphosphate farnesyltransferase (FDFT1), farnesyl diphosphate synthase (FDPS), hydroxymethylglutaryl CoA synthase (HMGCS1), isopentenyl-diphosphate δ-isomerase 1 (IDI1), lanosterol synthase (LSS), diphosphomevalonate decarboxylase (MVD), mevalonate kinase (MVK), squalene monooxygenase (SQLE) and transmembrane 7 superfamily member 2 (TM7SF2) was increased one- to two-fold in liver of T2D compared to CTRL (Figure 5b).

**Figure 5:**
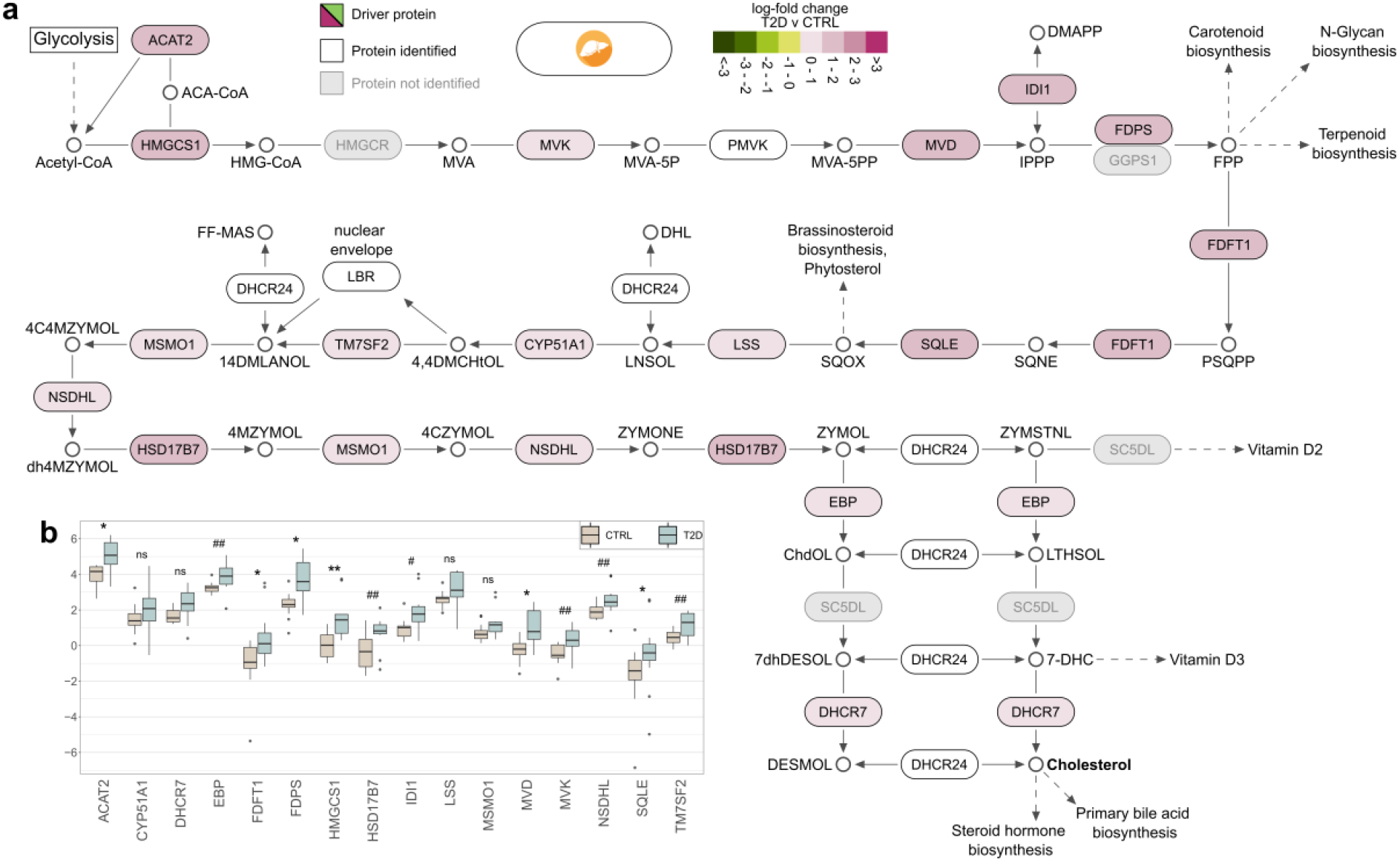
Detail on the increase of the biological pathway of cholesterol biosynthesis in liver. **a)** The pathway is based on KEGG and complemented with details from Reactome and the literature. Tiles illustrate log-fold change values of T2D compared to CTRL in liver. Proteins leading the increase of the pathway are marked as driving and are shown in black font and colored tiles. Proteins participating in the pathway and identified from MS proteomics, but not marked as leading edges are in white tiles, while those in grey tiles were not identified in our analysis. **b)** Detailed comparison of the abundance of driving proteins between CTRL and T2D in liver.

The cholesterol biosynthesis pathway initiates with acetyl-CoA that is produced by glycolysis and results in, among others, cholesterol that is utilized for the biosynthesis of steroid hormones and primary bile acids. The MS analysis identified the vast majority of proteins from key steps of the cholesterol biosynthesis pathway (Figure 5a). A detailed exploration on the protein abundancies in liver revealed that despite the moderate significance of the differences between CTRL and T2D in protein levels, their cumulative impact to the pathway was sufficient to lead to a very strong increase (Figure 5b; Supplementary Figure S16).

### Hemostasis is elevated in pancreatic islets of PD

The blood clot formation pathway of hemostasis was strongly enriched in pancreatic islets of PD (q<3.5×10^−6^). Specifically, the enrichment of the clotting cascade (q<8.6×10^−14^) was led by the upregulation of the common and intrinsic pathways that are responsible for fibrin clot formations (q<1.1×10^−6^ and q<4.5×10^−15^, respectively) (Figure 3; Supplementary Table S9). A large collection of 44 proteins that were higher in PD than in CTRL drove the identification of the significant increase of the pathway (Supplementary Figure S17a). Visualization of a subset of 21 proteins from the pathway of hemostasis illustrated the key biological processes affected by these enzymes (Figure 6). We observed a greater than 3-fold increase in transferrin (TF) and a solid elevation in multiple members of the serpin family of proteins (Figure 6; Supplementary Figure S17). Additionally, we observed that the significant increase of the 44 proteins associated with hemostasis was followed by a significant decrease in T2D that, in turn, was significantly higher than CTRL (Figure 3; Supplementary Figure S17b). In contrast to islets, hemostasis-related pathways in liver were significantly lower in T2D than in PD (Figure 3).

**Figure 6:**
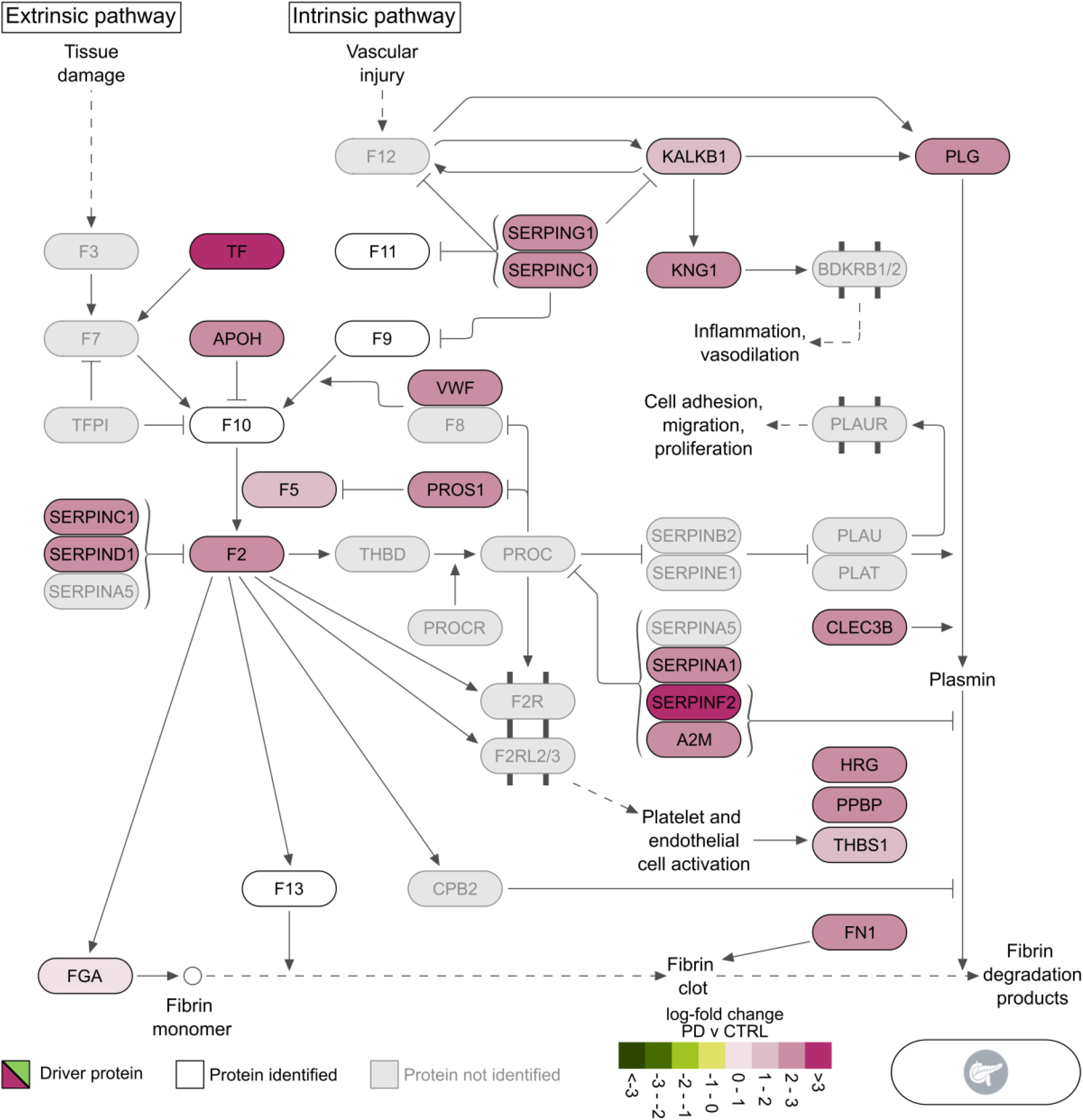
Biological pathway of hemostasis (coagulation cascade). The pathway is based on KEGG and complemented with details from Reactome and the literature. Tiles illustrate log-fold change values of PD compared to CTRL in pancreatic islets. Proteins leading the increase of the pathway are marked as driving and are shown in black font and colored tiles. Proteins participating in the pathway and identified from MS proteomics, but not marked as leading edges are in white tiles, while those in grey tiles were not identified in our analysis.

### Complement cascade increases in liver and pancreatic islets of PD

The complement cascade was significantly higher in PD than in CTRL in liver and pancreatic islets (q<8.4×10^−5^ and q<1.6×10^−72^, respectively) (Figure 3; Supplementary Table S9). Exploration of the details in the pathway revealed that the classical and lectin pathways, that are responsible for the creation of the C2 and C4 activators, were strongly elevated in pancreatic islets (q<2.8×10^−53^), but did not cross the threshold of statistical significance in liver (Figure 3; Supplementary Table S9). The upregulation of the pathway in liver was led by 20 proteins, while the one in pancreatic islets by 34 proteins, of which 14 were shared (Figure 7; Supplementary Figures S18a and S18b). The complement cascade remained higher in T2D than CTRL in pancreatic islets in contrast to liver where it significantly decreased (Figure 3). The two tissues were in agreement again when comparing T2D with PD. Specifically, the levels of the driving proteins for the upregulation of the complement cascade in PD decreased strongly in T2D resulting in an overall downregulation of the pathway from PD to T2D (q_liver_<1.4×10^−14^ and q_islets_<1.1×10^−16^) (Figure 3; Supplementary Figures S18c and S18d). The latter clearly marked an abrupt increase of immune system-related proteins in PD followed by a strong decrease in T2D that in liver leads to significantly higher levels in T2D compared to CTRL (Supplementary Figures S18c and S18d).

**Figure 7:**
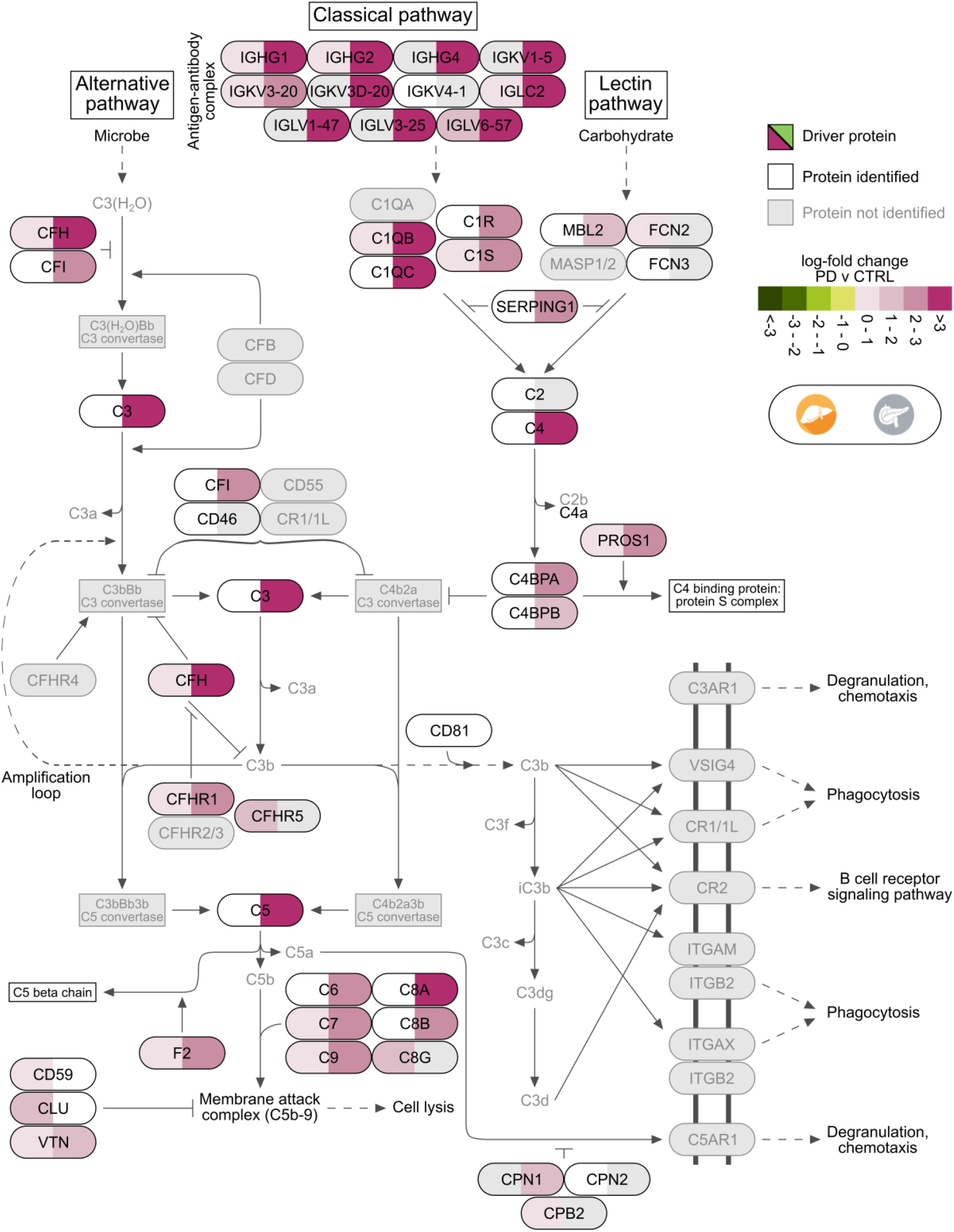
Biological pathway of complement cascade. The pathway is based on KEGG and complemented with details from Reactome and the literature. Tiles illustrate log-fold change values of PD compared to CTRL in liver (left hand side of tile) and pancreatic islets (right hand side of tile). Proteins leading the increase of the pathway are marked as driving and are shown in black font and colored tiles. Proteins participating in the pathway and identified from MS proteomics, but not marked as leading edges are in white tiles, while those in grey tiles were not identified in our analysis.

## Discussion

We performed a mass spectrometry-based proteomics analysis of samples from VAT, liver, skeletal muscle, pancreatic islets and serum from a cohort of 43 multi-organ donors (Figure 1). To our knowledge, this is the first proteomics study of this scale investigating PD and T2D across all key metabolic tissues in humans. Importantly, we provided the deepest proteome coverage to date for pancreatic islets. We performed a multi-dimensional exploratory analysis that highlighted biological processes and pathways altered along with the development of T2D (Figures 2 & 3). This map of defects in pathways was complemented with a fine-grained collection of proteins that were shown to drive their direction of change, as well as, their magnitude (Figures 4-7). Our findings highlighted several novel and well-characterized biological pathways to be significantly altered in PD and T2D, and, most importantly, paved the way for further intra-individual investigations. Overall, our study provides a unique resource to the field of metabolic diseases.

The vast majority of the biological processes that were increased in PD and T2D of pancreatic islets and liver were related to immune responses and their regulation (Supplementary Figure S10a). Links between the activation of the adaptive and the innate immune system and the progression of T2D have been demonstrated in earlier studies (de Candia et al., 2019; Itariu and Stulnig, 2014). However, the most interesting group in the PD network was the upregulation of biological processes involved in the regulation of the immune response in pancreatic islets and liver (Figure 2a). Liver also showed divergent patterns of alterations in PD and T2D, while in VAT and skeletal muscle there was a prevalent decrease in T2D (Figures 2a & 2b; Supplementary Figure S10). A comparison between T2D and PD not only provided insights for the tissues with strong responses to overt T2D, but also underlined significantly affected biological processes (Figure 2c). We observed no enriched biological processes or pathways in PD of VAT and skeletal muscle, while we identified very few differences between PD and T2D in pancreatic islets. In general, pancreatic islets dominated the alterations between PD-CTRL, and while proceeding chronologically towards T2D, they did not demonstrate further notable alterations. Unlike pancreatic islets, in liver there were over 150 enriched biological processes mainly related to the immune system, that differed between PD and T2D. In summary, our data suggests that the majority of the biological alterations in PD occur primarily in pancreatic islets and secondarily in liver, and during the progression and the establishment of overt T2D, alterations related to oxidative phosphorylation and cellular respiration appear in VAT and skeletal muscle (Figure 2).

The TCA cycle has previously been suggested to be a marker of insulin resistance in various tissues that we, in agreement with other studies, find to be strongly downregulated in T2D of skeletal muscle and VAT (Figures 3 & 4) (Befroy et al., 2007; Gaster, 2009; Heinonen et al., 2015). The amphibolic nature of the pathway involves the synthesis of fatty acids and amino acids, besides driving synthesis of ATP from ADP. In other words, its overall downregulation will affect the ATP production and other important metabolic pathways (Gaster et al., 2012). The pivotal role of mitochondrial RET in T2D was highlighted in a recent study that successfully treated T2D in mice models using magnetic and electric fields (Carter et al., 2020). The impairment of the TCA cycle and the RET chain in VAT and skeletal muscle of T2D subjects compared to CTRL and PD, in combination with the indistinguishable differences between CTRL and PD suggested a relatively late event becoming evident when diabetes is already established, rather than during its development (Figure 3). Taken together our data suggested the tissues and the T2D stage at which the downregulation of the TCA cycle and the RET chain become prominent.

A similar sequence of events applied to the enhanced cholesterol biosynthesis in liver. This may be driven by the enhanced transcription activation of genes encoding various key-enzymes of the pathway and may be linked to subsequent dysregulation of cholesterol and lipoprotein production (Figures 3 & 5). This also became evident in overt T2D and is likely to be a consequence rather than a cause of the disease. The elevated levels of many enzymes of cholesterol biosynthesis was likely driven by two biological pathways that have SREBP as their core regulator, namely regulation of cholesterol biosynthesis by SREBP and gene expression activation by SREBP (Figure 3). We hypothesize that it is driven by increased levels and activity of SREBP transcription factors. The latter may subsequently lead to increased synthesis or assembly of other lipid moieties, such as triacylglycerols, and it may favor the production and release into the circulation of lipoproteins, such as VLDL particles (Brown and Goldstein, 1997; Shimano and Sato, 2017).

Liver, skeletal muscle and VAT are the classical sites for insulin-regulated substrate metabolism, accounting for much of energy expenditure and storage (Hong et al., 2014; Viscarra and Ortiz, 2013). We identified patterns of enhanced lipid, in particular cholesterol, synthesis in the liver and reduced mitochondrial function, and hence glucose and lipid oxidation, in VAT and skeletal muscle (Figure 3). Taken together these alterations are compatible with less utilization of energy substrates in muscle and adipose, redistribution of lipids towards the liver and altered lipoprotein release. It appears that these alterations in energy metabolism occur later than the islet perturbations, and, in fact, they can partly be secondary to a relative insulin deficiency. Nonetheless, the alterations in liver, muscle and VAT are certainly important in maintaining the diabetic state including hyperglycemia, dyslipidemia and overweight. The established endocrine dysfunction of the pancreas as well as adaptative mechanisms of the brain are also involved in ‘defending’ chronic hyperglycemia in T2D (Alonge et al., 2021; DeFronzo, 2009; Lundqvist et al., 2020).

IR with compensatory hyperinsulinemia are common features of T2D and impaired glucose tolerance, that are subsequently associated with increased risk of coronary heart disease (CHD) (Haffner Steven M. and Hanley Anthony J.G., 2002). A possible tie between hyperinsulinemia and CHD can be found in the hypercoagulable state that is characteristic of patients with T2D (Yamada et al., 2000). Several studies have indicated a connection between hyperinsulinemia/hyperglycemia and activation of the coagulation cascade following food intake (Ceriello et al., 1996, 1995). These findings have been further supported by reports that infusion of glucose induces a transient increase in the generation of thrombin (F2) in normal subjects, and of the effect being more pronounced in T2D patients. Here, we observed strong elevation in the levels of TF, F2 and several members of the serpin family that play important roles in the formation of fibrin clots and plasmin (Figure 6). Moreover, multiple serine proteases of the coagulation system have been shown to function as alternative activators of C5 and C3 of the complement cascade (Amara et al., 2008). Both pathways were strongly upregulated in PD of pancreatic islets while the latter was also upregulated in liver of PD (Figures 3, 6 & 7).

It is of great interest that pancreatic islets, in contrast to liver, VAT and skeletal muscle, displayed strong perturbations of protein patterns in PD individuals compared to CTRL (Figures 2 & 3). Evidence for increased hemostatic and inflammatory activity in PD was not further enhanced in T2D subjects (Figure 3). Instead, they retained similar patterns to PD for immune responses, while there was even a partial return towards the levels of CTRL with respect to complement activation and hemostasis via platelet and coagulation activation (Supplementary Figures S17 & S18). In conclusion, this data is compatible with an early, potentially causal role, of vascular, inflammatory and immune impairment within the pancreatic islets that lead to dysfunction of their endocrine cells, that may be manifested by insufficient insulin production and increased glucagon levels (D’Alessio, 2011; DeFronzo, 2009). It could therefore be speculated that there is a vascular/inflammatory insulitis to pancreatic islets that occurs mainly during diabetes development but later on subsides.

In this study we were unable to infer causality from the results due to the limited number of subjects that in turn leads to reduced statistical power. Our results would need to be cross validated in other settings involving additional experimental procedures to be further confirmed and, potentially, explored in medical settings. We were restricted only to the reported clinical variables due to the Swedish legislation on obtaining details for other underlying diseases and medication that the subjects were administered prior to their admission to the emergency and ICU at the hospital. Moreover, in our analysis we have adjusted for potential confounding technical factors including CIT and the purity of the samples of pancreatic islets. However, sampling methods at the different medical facilities and the deep-frozen storage of the samples may have impacted the results to some extent.

In conclusion, we reported an extensive collection of biological pathways that were found to be altered across several metabolically relevant tissues in the development of type 2 diabetes. More importantly, we presented the first multi-tissue proteomics map describing the chronological order of tissue-specific metabolic dysregulation across healthy, prediabetes and type 2 diabetes subjects.

## Methods

### Ethics declaration

The collection and utilization of human organs for scientific research and transplantation purposes is regulated by the Swedish legislation. The human tissue lab is a biobank for multi-organ donors funded by the excellence of diabetes research in Sweden (EXODIAB), and is a collaborative initiative between the universities of Uppsala and Lund. Samples (n = 43) for five metabolically-relevant tissues were obtained from the human tissue lab following the Uppsala Regional Ethics Committee approval (Dnr: 2014/391). Informed consent was received from the donors or their legal guardians for their organs to be used for scientific research. Storage and analysis of the samples has been in full accordance with the Swedish law and regional standard practices. No tissue samples were obtained from prisoners.

### Sample collection

Frozen samples of VAT, liver, skeletal muscle, pancreatic islets and serum were obtained for 43 multi-organ donors. The world health organization guidelines were followed to identify T2D subjects in the corresponding medical facilities (World Health Organization, 2016). PD was identified based on the percentage of glycosylated hemoglobin in blood (5.7%≤HbA_1c_≤6.5%), while normoglycemia (CTRL) was assigned otherwise (HbA_1c_≤5.6%) (Supplementary Table S1). Glucose-stimulated insulin secretion (GSIS) and purity of the pancreatic islet samples were measured (Krogvold et al., 2015). Information on characteristics of the anthropometric, technical, T2D-related and ICU-related variables was available for the study and was described in (Supplementary Table S1). We also explored merging PD with T2D and comparing to CTRL, additionally to merging PD with CTRL and comparing to T2D.

### Mass spectrometry proteomics

#### Proteomic sample preparation

Skeletal muscle, VAT and liver samples were lysed in Sodium Deoxycholate containing lysis buffer (1% Sodium Deoxycholate (SDC), 10 mM Tris (2-carboxyethyl) phosphine (TCEP), 40 mM Chloroacetamide (CAA) and 100 mM of Tris (pH 8.5)). The samples were homogenized with an Ultra Turrax homogenizer (IKA). In case of the pancreatic islets extract, lysis buffer containing 10% 2,2,2-trifluoroethanol (TFE), 10 mM Tris (2-carboxyethyl) phosphine (TCEP), 40 mM Chloroacetamide (CAA) and 100 mM of Tris (pH 8.5) was used for protein extraction. For the serum samples, 1µl of serum was mixed with 24 µl of 1% SDC buffer. Samples were boiled at 95°C for 10 min. Tissue samples were sonicated using a tip sonicator while serum samples were sonicated using water bath sonicator. Tissue lysates were then centrifuged at 16000g for 10 min and the supernatant was collected for protein digestion. Proteins were digested using the endoproteinases LysC and trypsin (1:100 w/w) at 37°C overnight with shaking. Digested peptides were acidified using 1% Trifluoroacetic acid (TFA) and purified using the StageTips containing SDB-RPS material and eluted in 1% ammonia and 80% acetonitrile. Peptides were concentrated, dried using Speed-vac and res-suspended in buffer containing 2% acetonitrile and 0.1% TFA.

#### Peptide libraries, high pH reversed-phase fractionation

For serum, skeletal muscle, VAT and liver we used peptide libraries generated in house for quantification based on data-independent acquisition (DIA). To generate the library for the pancreatic islets, 15 µg of peptides were pooled from few samples and fractionated using high pH reversed-phase chromatography (Kulak et al., 2017). 16 fractions were automatically concatenated using a rotor valve shift of 90 s. Approximately 0.3 µg of each fraction were subjected to LC-MS/MS measurements via data-dependent acquisition (DDA).

#### Mass spectrometry

Peptides were measured using LC-MS instrumentation consisting of an Easy nanoflow HPLC system (Thermo Fisher Scientific, Bremen, Germany) coupled via a nanoelectrospray ion source (Thermo Fischer Scientific, Bremen, Germany) to a Q Exactive HF-X mass spectrometer. Purified peptides were separated on a 50 cm C18 column (inner diameter 75 µm, 1.8 µm beads, Dr. Maisch GmbH, Germany). Peptides from the tissue samples were loaded onto the column with buffer A (0.5% formic acid) and eluted with a 100 min linear gradient increasing from 2-40% buffer B (80% acetonitrile, 0.5% formic acid). For the peptides from serum samples 45 min linear gradient was used. After the gradient, the column was washed with 90% buffer B and re-equilibrated with buffer A.

Mass spectra were acquired in either DDA or DIA mode. For the pilot experiment where samples from six subjects were analyzed, data for tissue samples was obtained in DDA while the serum data was acquired in DIA mode. For the main experiments, data for all samples was acquired in DIA mode. For the pancreatic islets extract, the library samples were measured in DDA mode. For the DDA method, the mass spectra were acquired with automatic switching between MS and MS/MS using a top 15 method. MS spectra were acquired in the Orbitrap analyzer with a mass range of 300-1750 m/z and 60,000 resolutions at m/z 200 with a target of 3 × 10^6^ ions and a maximum injection time of 25 ms. HCD peptide fragments acquired at 27 normalized collision energy were analyzed at 15000 resolution in the Orbitrap analyzer with a target of 1 x 105 ions and a maximum injection time of 28 ms. A DIA MS method was used for all tissue proteome measurements in which one full scan (300 to 1650 *m/z*, resolution = 60,000 at 200 *m/z*) at a target of 3 × 10^6^ ions was first performed, followed by 32 windows with a resolution of 30000 where precursor ions were fragmented with higher-energy collisional dissociation (stepped collision energy 25%, 27.5%, 30%) and analyzed with an AGC target of 3 × 10^6^ ions and maximum injection time at 54 ms in profile mode using positive polarity. For the serum samples, a DIA MS method in which one full scan (300 to 1650 *m/z*, resolution = 120,000 at 200 *m/z*) at a target of 3 × 10^6^ ions was first performed, followed by 22 windows with a resolution of 30000 where precursor ions were fragmented with higher-energy collisional dissociation (stepped collision energy 25%, 27.5%, 30%) and analyzed with an AGC target of 3 × 10^6^ ions and maximum injection time at 54 ms in profile mode using positive polarity.

#### Data processing

Raw MS files from the experiments measured in the DDA mode (pancreatic islets library & tissue samples from pilot experiments) were processed using MaxQuant (Cox and Mann, 2008). MS/MS spectra were searched by the Andromeda search engine (integrated into MaxQuant) against the decoy UniProt-human database (downloaded in December 2017) with forward and reverse sequences. In the main Andromeda search precursor, mass and fragment mass were matched with an initial mass tolerance of 6 ppm and 20 ppm, respectively. The search included variable modifications of methionine oxidation and N-terminal acetylation and fixed modification of carbamidomethyl cysteine. The false discovery rate (FDR) was estimated for peptides and proteins individually using a target-decoy approach allowing a maximum of 1% false identifications from a revered sequence database. Raw files acquired in the DIA mode were processed using Biognosys Spectronaut software version 13 (Bruderer et al., 2015). A single peptide library was generated in Spectronaut using the combined MaxQuant search results for the DDA runs from the pancreatic islets sample. The experimental DIA runs were then analyzed in Spectronaut using default settings.

### Computational analysis

We used precompiled gmt files containing collections of GO terms (C5) and biological pathways for KEGG and Reactome (C2) from the molecular signatures database (MSigDB) (Supplementary Table S11) (Jassal et al., 2020; Kanehisa et al., 2017; Liberzon et al., 2011; Subramanian et al., 2005). The collection of gmt files was initially used to compile a universe of unique gene names. For each quantified protein, the DIA provides an ordered list of UniProt identifiers and gene names ranked based on statistical confidence. The universe was used as means to decide on the unique gene names that most accurately represent the quantified proteins. If one or more identifiers from the ordered list intersected with the universe then the first hit from the ordered list was selected, otherwise the first, and most significant, of the three identifiers was used.

Raw data was log_2_-transformed to better approximate a normal distribution and proteins identified in at least 80% of the samples were retained for downstream analysis (Supplementary Figure S4a). Abundancies of proteins corresponding to the same gene in each tissue were averaged within samplea. Samples with consistent large deviations from the median abundancies of all samples across tissues were excluded from the analysis (Supplementary Figure S3; Supplementary Table S2) and normalization was performed using median sweeping (Supplementary Figure S4b). Missing protein levels were strongly biased for proteins located on the lower end of the detection limit (Supplementary Figure S5a & S5b); hence, in order to impute the missing values we selected a method that draws random values from a truncated distribution with parameters estimated from quantile regression (QRLIC) using the function *impute*.*QRILC* from the R package *imputeLCMD* (Supplementary Figure S5c & S5d).

#### Differential analysis

The differential analysis was performed on pairwise comparisons among CTRL, PD and T2D, and the merged groups CTRL+PD and PD+T2D using the R package *limma* (Ritchie et al., 2015). The differential models were corrected for confounding factors explaining ≥1% of the median variance across proteins in at least one tissue (Supplementary Figure S1; Supplementary Table S4) (Hoffman and Schadt, 2016). Raw p-values were FDR-corrected to q-values, and they were subsequently combined with the log-fold change (logFC) to compute π-values: π=logFC × -log_10_(q-value) (Storey, 2003; Xiao et al., 2014).

#### GO and pathway enrichment analysis

Lists of proteins ranked on π-values from the differential analysis were imported into the functions *cameraPR* and *fgsea* from the R packages *limma* and *fgsea*, respectively (Korotkevich et al., 2019; Wu and Smyth, 2012). *cameraPR* was executed on default settings, while *fgsea* was set to perform 1M permutations, and to consider only terms with ≥5 and ≤300 proteins. For both methods the enrichment of GO terms and biological pathways was performed separately using the *gmt* files described above and the raw p-values were corrected for multiple testing using the R package *qvalue*. As the collection of enriched GO terms and pathways the intersection of the significant terms (q<0.05) sharing the same direction of change in both methods was considered.

Enrichment analysis of biological terms for pre-defined sets of proteins identified only in one tissue (tissue-specific) and shared among tissues (tissue-shared) was performed using a hypergeometric test. The background set for tissue-shared proteins was the union of proteins identified across tissues, while as background set for tissue-specific proteins was used the collection of proteins in the respective tissue. Sets with ≥5 and ≤300 proteins were retained and p-values were corrected for multiple testing using the R package *qvalue*.

## Data and code availability

Raw MS proteomics files are available upon request and will be available at the time of review in the PRoteomics IDEntification Database (PRIDE). Restriction of access applies to anthropometric, clinical and technical variables. This information can be available to researchers whose ethical permit applications to EXODIAB meet the criteria to allow confidential data access. The pipeline for the analysis is available on GitHub *https://github.com/klevdiamanti/multitissue_ms_proteomics* (Supplementary Table S12). A web-based tool that allows visualization and exploration for levels of proteins across tissues is available on *http://bioinf.icm.uu.se:3838/multitissue_ms_proteomics/*.

## Author contributions

KD analyzed the data, interpreted the results and wrote the first draft of the manuscript. MC prepared samples, prepared curated groups of GO terms and assisted in writing the manuscript. GP prepared samples. MJP, JWE, OK and CLL interpreted the results and assisted in writing the manuscript. CK provided comments on the analysis. MM, FM and ASD performed MS measurements and provided comments on the analysis and the manuscript. JK provided the infrastructure for the web-based exploratory tool, contributed to the initial study design and comments on the manuscript. CW designed and supervised the study, and assisted in writing the manuscript.

## Acknowledgements

Funding was provided by AstraZeneca to CW; European Commission Marie Skłodowska Curie Innovative Training Network TREATMENT (H2020-MSCA-ITN-721236) to JWE and MJP; the Swedish Diabetes Foundation (Swedish Diabetes Association) to CW, JWE, OK and MJP; the Ernfors foundation to CW, JWE, OK and MJP; the Excellence for Diabetes Research in Sweden (EXODIAB) to CW, JWE, OK and MJP; the Zetterling Foundation to JWE and MJP; the Swedish Research Council to OK; the Novo Nordisk Foundation (NNF) to OK and JWE; the eSSENCE grant to JK; an unconditional donation from NNF to the NNF Center for Basic Metabolic Research (http://www.cbmr.ku.dk) (NNF18CC0034900) and the NNF Center for Protein Research (NNF14CC001). The authors would also like to acknowledge Rebeca Soria Romero and the mass spectrometry platform from the NNF Center for Protein Research for technical assistance and access to mass spectrometers.

