## Supplementary Figures for "Organ-specific metabolic pathways distinguish prediabetes, type 2 diabetes and normal tissues"

[1]Department of Immunology, Genetics and Pathology, Uppsala University, Uppsala, Sweden
[2]Department of Medical Sciences, Clinical Diabetes and Metabolism, Uppsala University, Uppsala, Sweden
[3]Translational Science & Experimental Medicine, Early Cardiovascular, Renal and Metabolism, BioPharmaceuticals R&D, AstraZeneca, Gothenburg, Sweden
[4]Karolinska Institutet/AstraZeneca Integrated CardioMetabolic Center (KI/AZ ICMC), Department of Medicine, Novum, Huddinge, Sweden
[5]Novo Nordisk Foundation Center for Protein Research, Faculty of Health Sciences, University of Copenhagen, Denmark
[6]Department of Oncology-Pathology, Karolinska Institute, Stockholm, Sweden
[7]Science for Life Laboratory, Department of Cell and Molecular Biology, Uppsala University, Uppsala, Sweden
[8]Institute of Computer Science, Polish Academy of Sciences, Warsaw, Poland
[9]Washington National Primate Research Center, Seattle, WA, USA
[10]Swedish Collegium for Advanced Study, Uppsala, Sweden
[11]Novo Nordisk Foundation Center for Basic Metabolic Research, Faculty of Health and Medical Science, University of Copenhagen, Copenhagen, Denmark
[12]Department of Proteomics and Signal Transduction, Max Planck Institute of Biochemistry, Martinsried, Germany
[13]Department of Medicine, University of Gothenburg, Gothenburg, Sweden

#### *Supplementary Material*

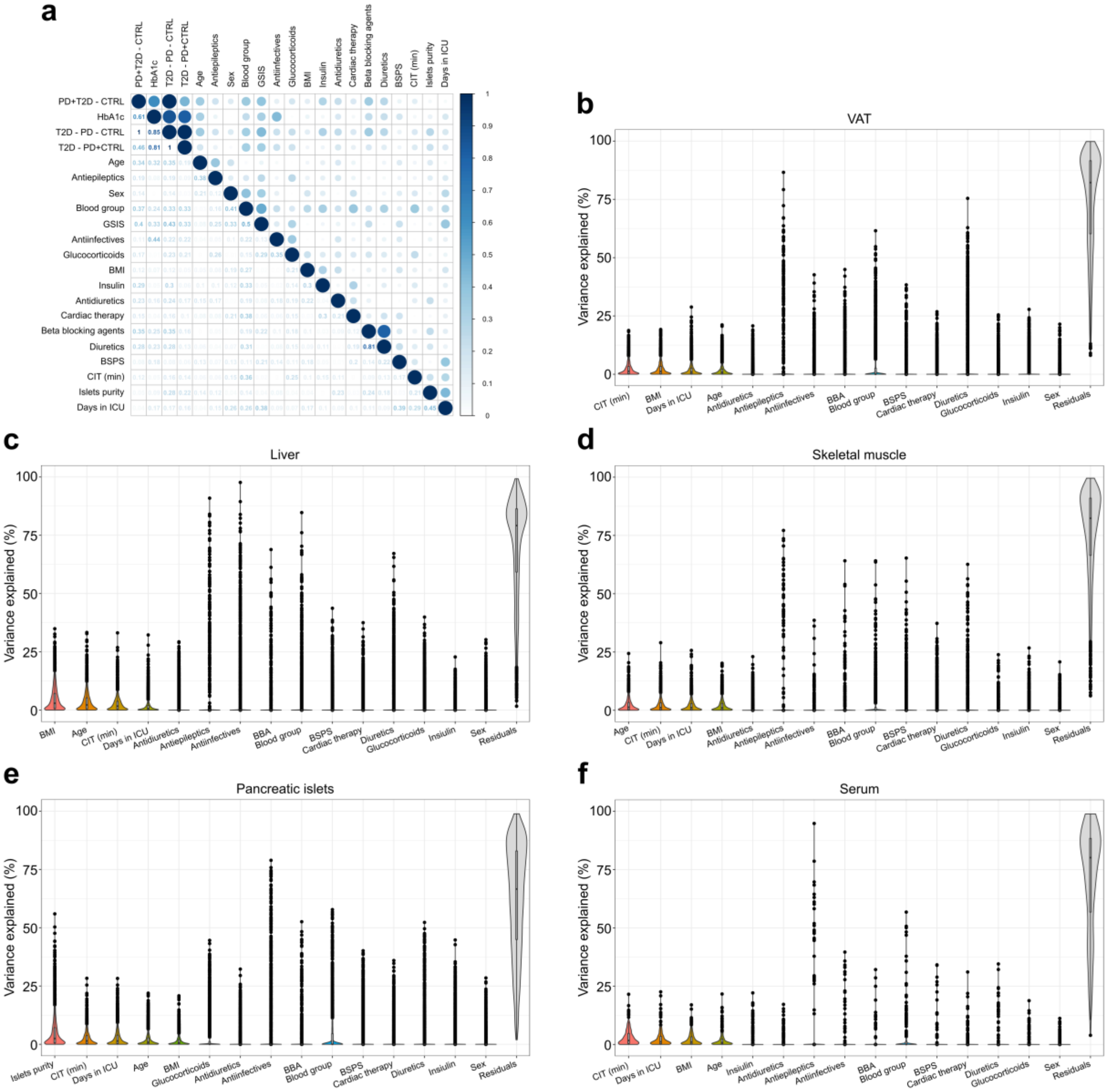


**Supplementary Figure S1: Correlation across clinical variables and the percentage of variation of the identified proteins they explain in each tissue. a)** Canonical correlation analysis across the set of clinical variables that includes anthropometric measurements (age, BMI, sex and blood group), technical variables (CIT and purity of islet sample), days hospitalized in the ICU and medication administered during this time (antidiuretics, antiepileptics, antiinfectives, beta blocking agents (BBA), blood substitutes and perfusion solutions (BSPS), cardiac therapy, diuretics, glucocorticoids and insulin). The analysis was performed using the function *canCorPairs* from the R package *variancePartition*. **b-f)** Violin plots represent percentage of variation of proteins within tissue explained by clinical variables. The detailed median and the standard deviation values for the percentage of variation are shown in (Supplementary Table S4) The analysis was performed using the function *fitExtractVarPartModel* from the R package *variancePartition*.

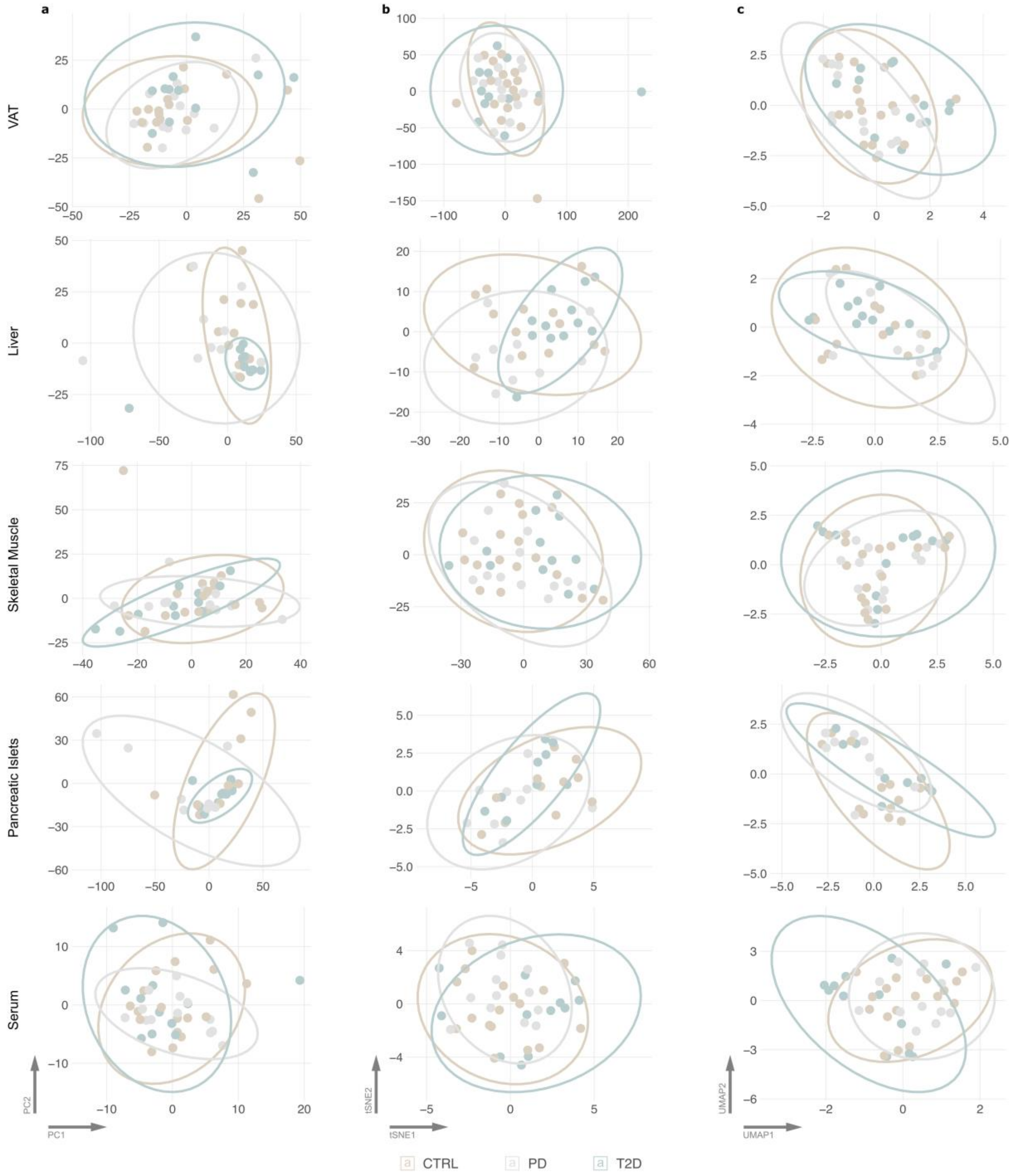


**Supplementary Figure S2: Representation of the protein abundancies for samples in a reduced dimensionality space. a)** The two first principal components (PCs) from the principal component analysis (PCA) that was performed using the *pca* function from the R package *pcaMethods* with the selected method being “ppca”. **b)** The two first components from the t-distributed stochastic neighbor embedding (tSNE) that was performed on the 10 first PCs from the PCA analysis from *a*. Perplexity for the tSNE algorithm was set to 1/3 of the total number of samples in each tissue. **c)** The two first components from the uniform manifold approximation and projection (UMAP) that was performed on the 10 first PCs from the PCA analysis from *a*. The number of neighbors for UMAP was set to five.

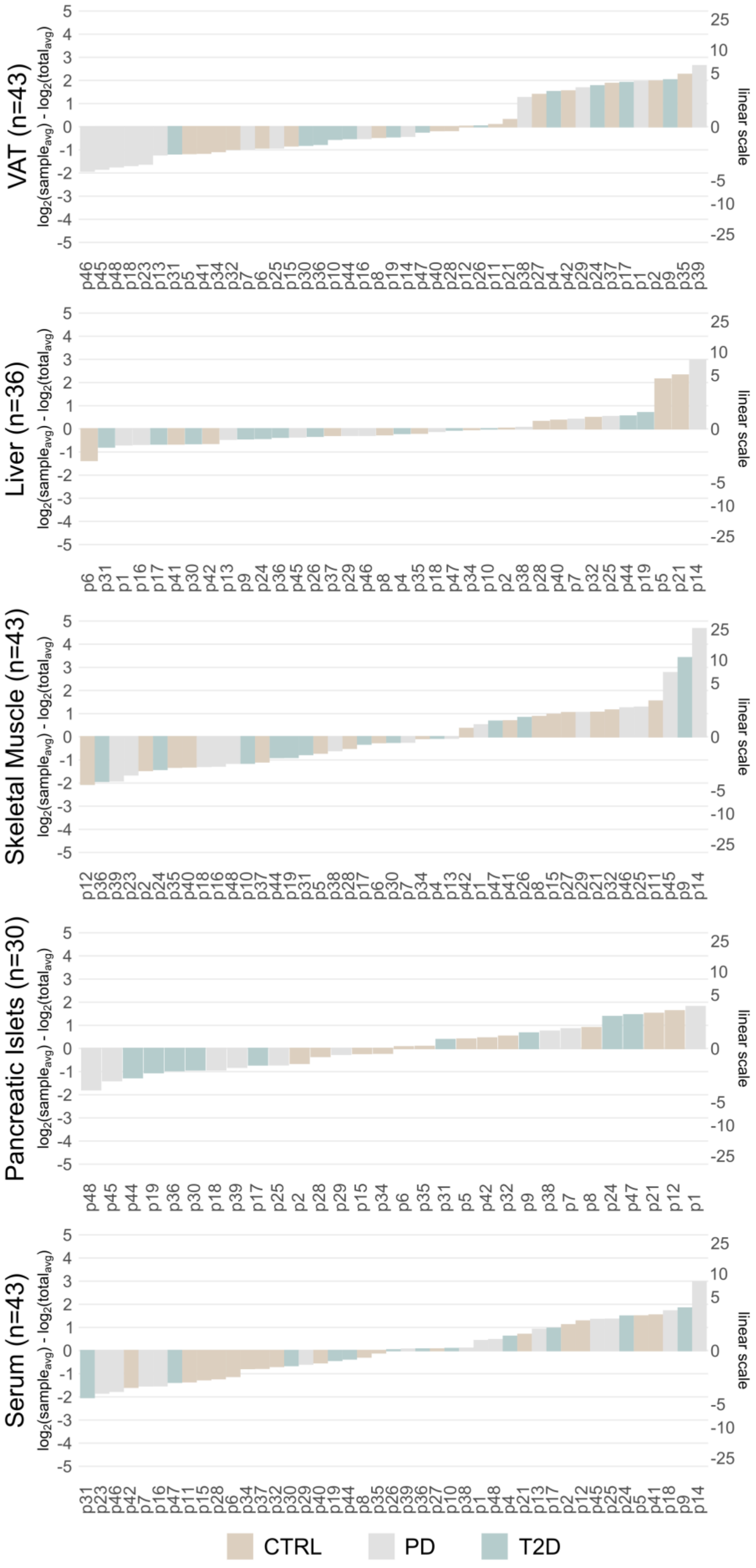
VAT (n=43)
Liver (n=36)
Skeletal Muscle (n=43)
Pancreatic Islets (n=30)
Serum (n=43)
$log_2(sample_{avg}) - log_2(total_{avg})$
linear scale
5
4
3
2
1
0
-1
-2
-3
-4
-5
25
10
5
0
-5
-10
-25
CTRL
PD
T2D

**Supplementary Figure S3: Difference of median protein abundance for samples compared to the average median abundance of all samples in the corresponding tissue.** Samples are shown in the x axis, while the left and right hand side y axes represent logarithmic and linear scales of the median differences, respectively.

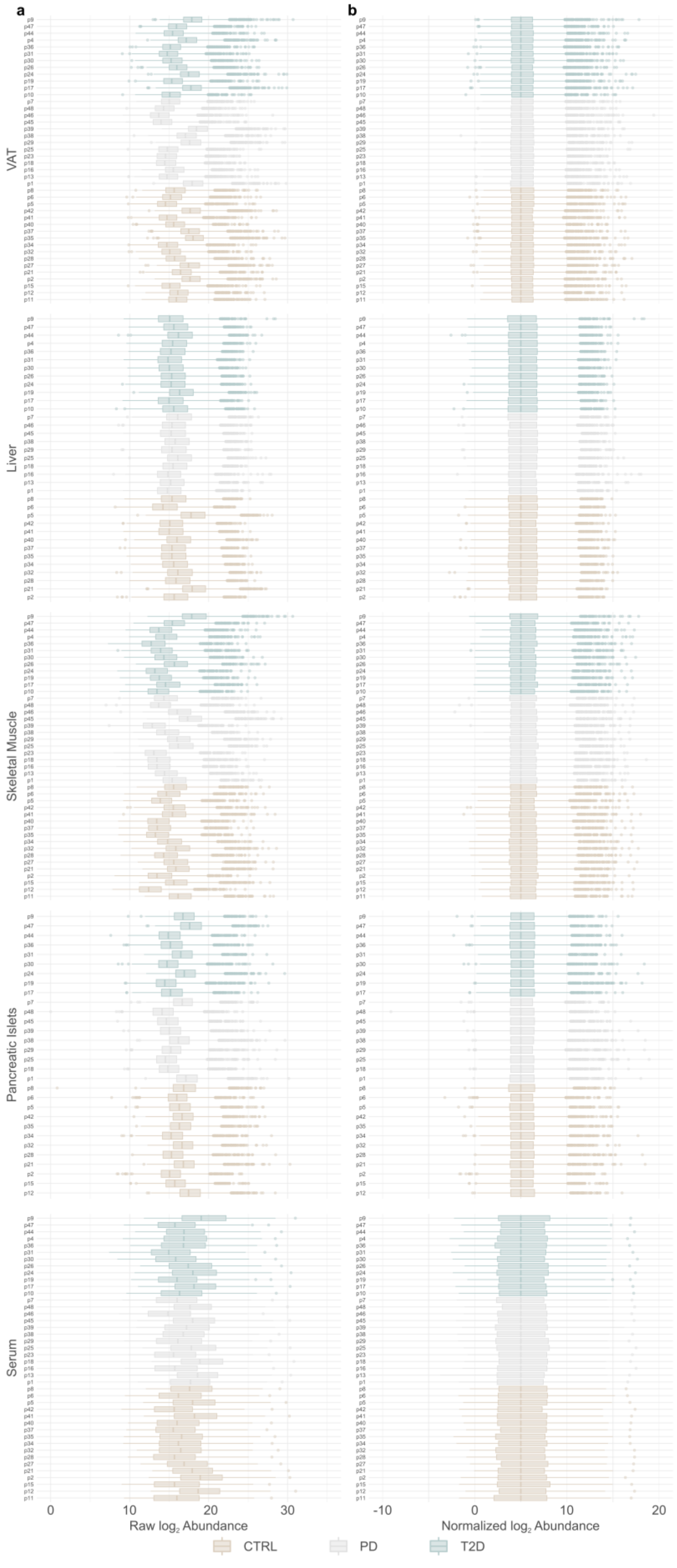
a
b
VAT
Liver
Skeletal Muscle
Pancreatic Islets
Serum
Raw log2 Abundance
Normalized log2 Abundance
CTRL
PD
T2D

**Supplementary Figure S4: Distribution of protein abundancies prior to and after normalization.** Sample IDs are shown in the y axis. **a)** Boxplots for raw $log_2$ protein levels. **b)** Boxplots for normalized protein levels.

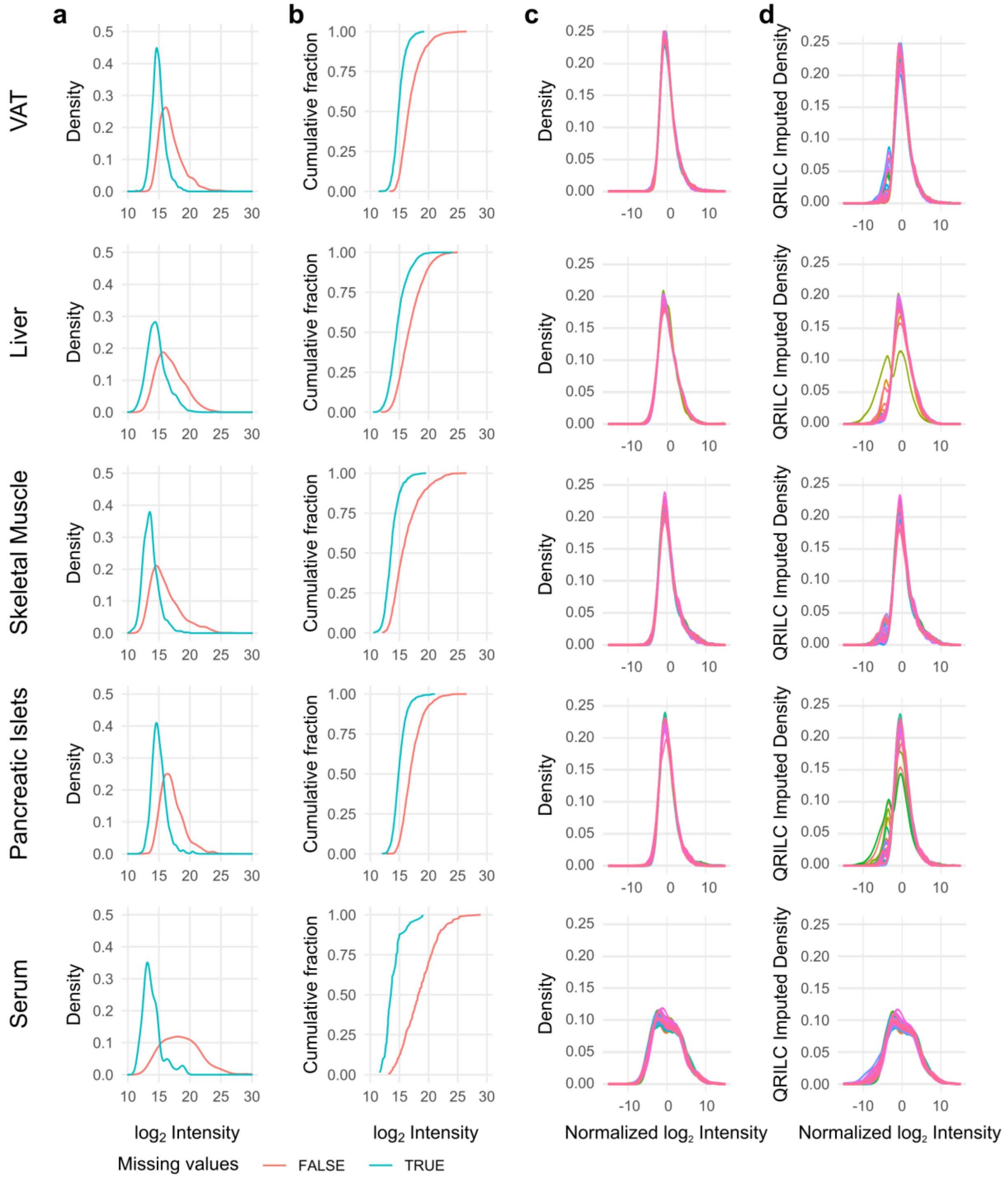


**Supplementary Figure S5: Summary of the distribution of protein abundancies prior and after imputation. a)** Distribution of the abundance of proteins with and without missing values. **b)** Cumulative fraction corresponding to *a*. **c)** Distribution of normalized $\log_2$ protein abundancies. **d)** Distribution of protein abundancies after QRILC lower-tail imputation.

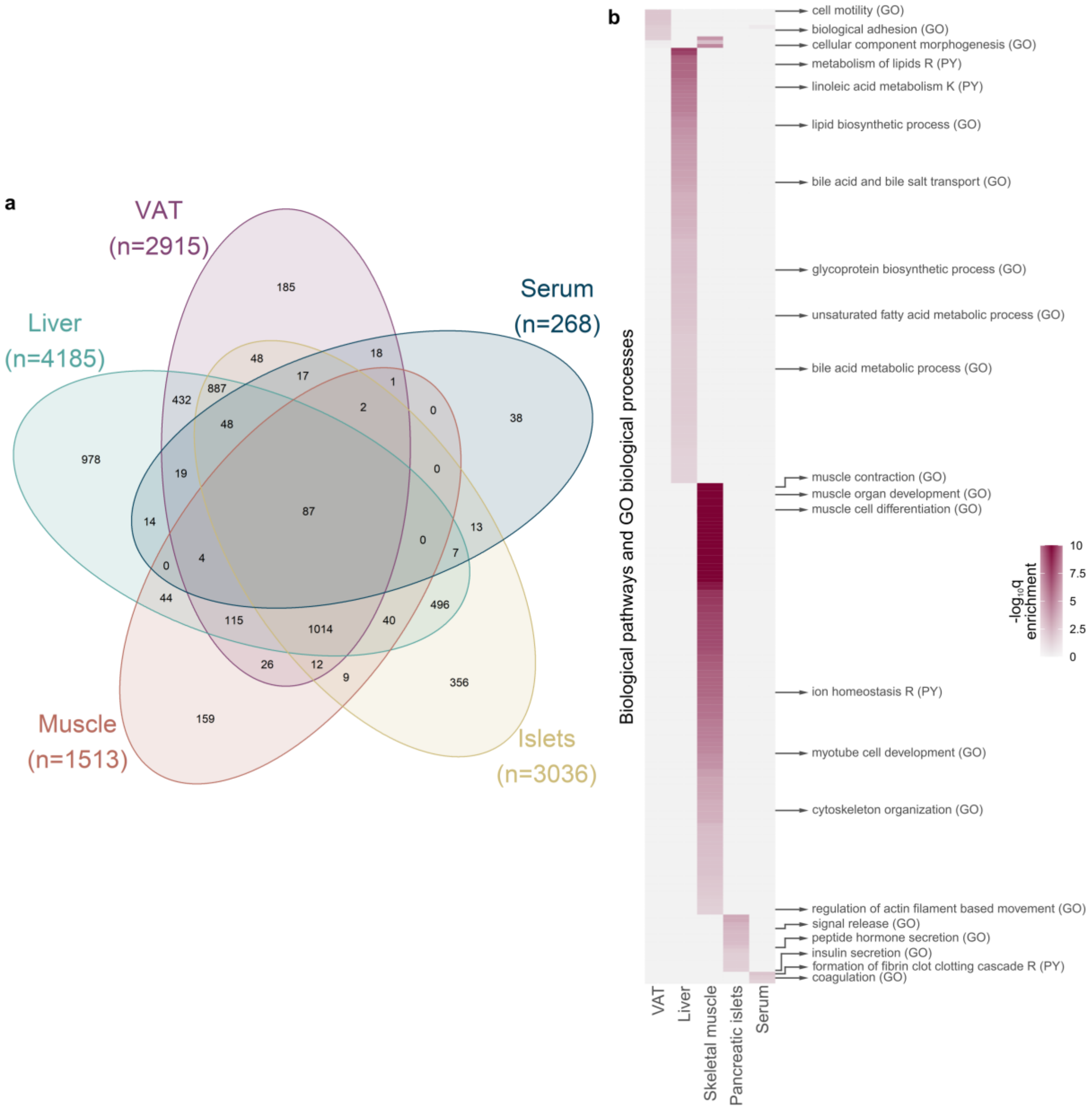


**Supplementary Figure S6: Exploration of tissue-specific proteins. a) Venn diagram of proteins shared among tissues and detected in single tissues. b)** Significantly enriched (q<0.05) GO terms for biological processes and biological pathways for proteins identified only in one tissue (tissue-specific). Columns represent tissues and rows represent enriched terms. Values in the parentheses of the terms mark the origin of the term: GO for gene ontology (GO) terms and PY for pathways. Terms representing pathways contain R or K as suffix to signify Reactome or KEGG pathways, respectively. A detailed list of the enriched terms is shown in (Supplementary Table S5).

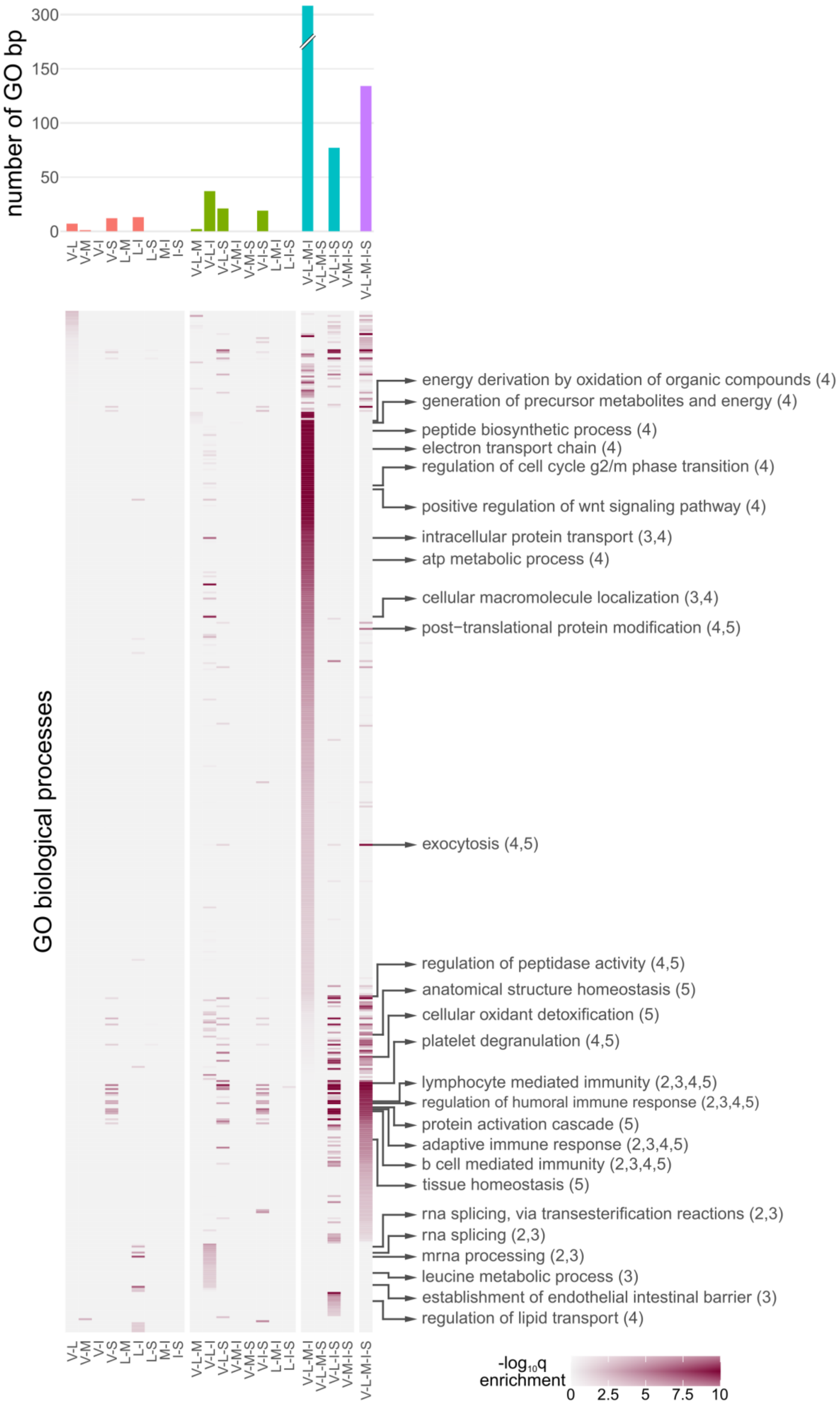
number of GO bp
300
150
100
50
0
V-L
V-M
V-I
V-S
L-M
L-I
L-S
M-I
I-S
V-L-M
V-L-I
V-L-S
V-M-I
V-M-S
V-I-S
L-M-I
L-I-S
V-L-M-I
V-L-M-S
V-L-I-S
V-M-I-S
V-L-M-I-S
GO biological processes
energy derivation by oxidation of organic compounds (4)
generation of precursor metabolites and energy (4)
peptide biosynthetic process (4)
electron transport chain (4)
regulation of cell cycle g2/m phase transition (4)
positive regulation of wnt signaling pathway (4)
intracellular protein transport (3,4)
atp metabolic process (4)
cellular macromolecule localization (3,4)
post−translational protein modification (4,5)
exocytosis (4,5)
regulation of peptidase activity (4,5)
anatomical structure homeostasis (5)
cellular oxidant detoxification (5)
platelet degranulation (4,5)
lymphocyte mediated immunity (2,3,4,5)
regulation of humoral immune response (2,3,4,5)
protein activation cascade (5)
adaptive immune response (2,3,4,5)
b cell mediated immunity (2,3,4,5)
tissue homeostasis (5)
rna splicing, via transesterification reactions (2,3)
rna splicing (2,3)
mrna processing (2,3)
leucine metabolic process (3)
establishment of endothelial intestinal barrier (3)
regulation of lipid transport (4)
-log10q
enrichment
0
2.5
5
7.5
10

**Supplementary Figure S7: Significantly enriched (q<0.05) GO terms for biological processes for proteins shared among tissues (tissue-shared).** The top panel shows the number of enriched terms for each combination of tissues. Panels from left to right represent proteins shared exclusively among two, three, four and five tissues. Columns represent tissue combinations and rows represent enriched GO terms. Column names are abbreviations of the tissue names: VAT (V), liver (L), skeletal muscle (M), pancreatic islets (I) and serum (S). Values in the parentheses of the GO terms mark the panel(s) in which the term is enriched. 2 implies combinations of 2 tissues, 3 combination of 3 tissues and so on. A detailed list of the enriched terms is shown in (Supplementary Table S6).

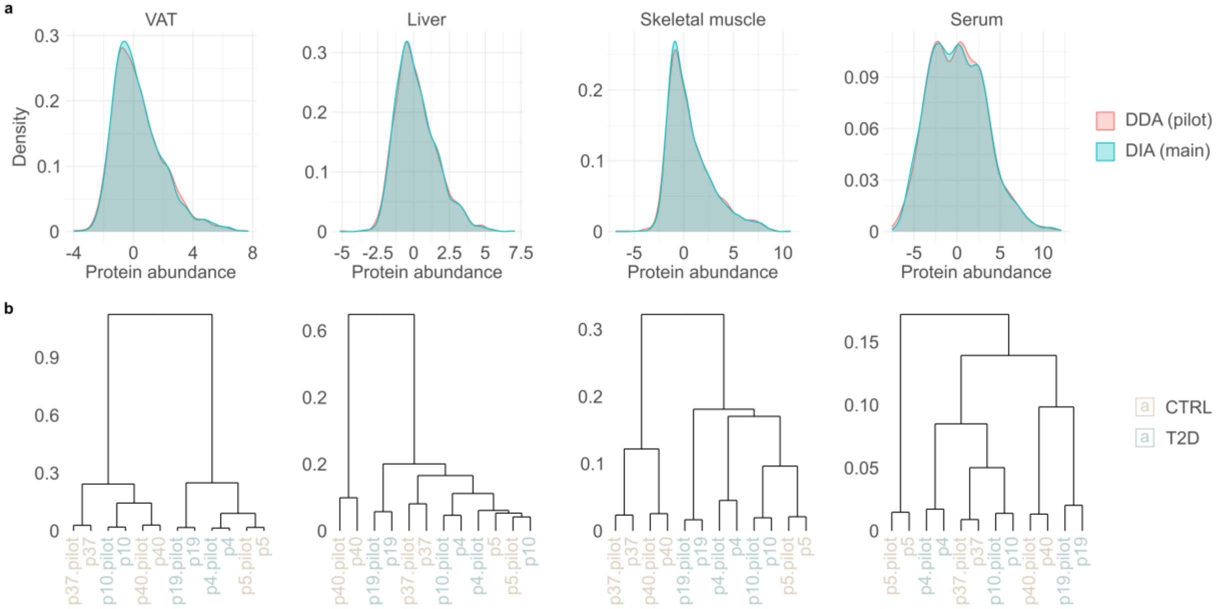


**Supplementary Figure S8: Comparison and clustering of six samples that were analyzed in both pilot and main analyses.** Pancreatic islets were excluded from the analysis due to low quality of four samples, which were consequently excluded from the main analysis. Samples included in this analysis are equally distributed between CTRL and T2D. Pilot study is marked as data-dependent acquisition (DDA) and the main one as data-independent acquisition (DIA). The data was $\log_2$-transformed and the set of shared top-500 most abundant proteins was analyzed. Batch effect was corrected using the function *removeBatchEffect* from the R package *limma*. **a)** Distribution of proteins in the pilot and main run after batch effect correction. **b)** Hierarchical clustering (HC) of DDA and DIA samples. DDA samples were marked with the suffix “pilot”. *1-r*, where *r* is Pearson correlation coefficient, was used as the distance metric and the *ward.D* algorithm was used for the HC.

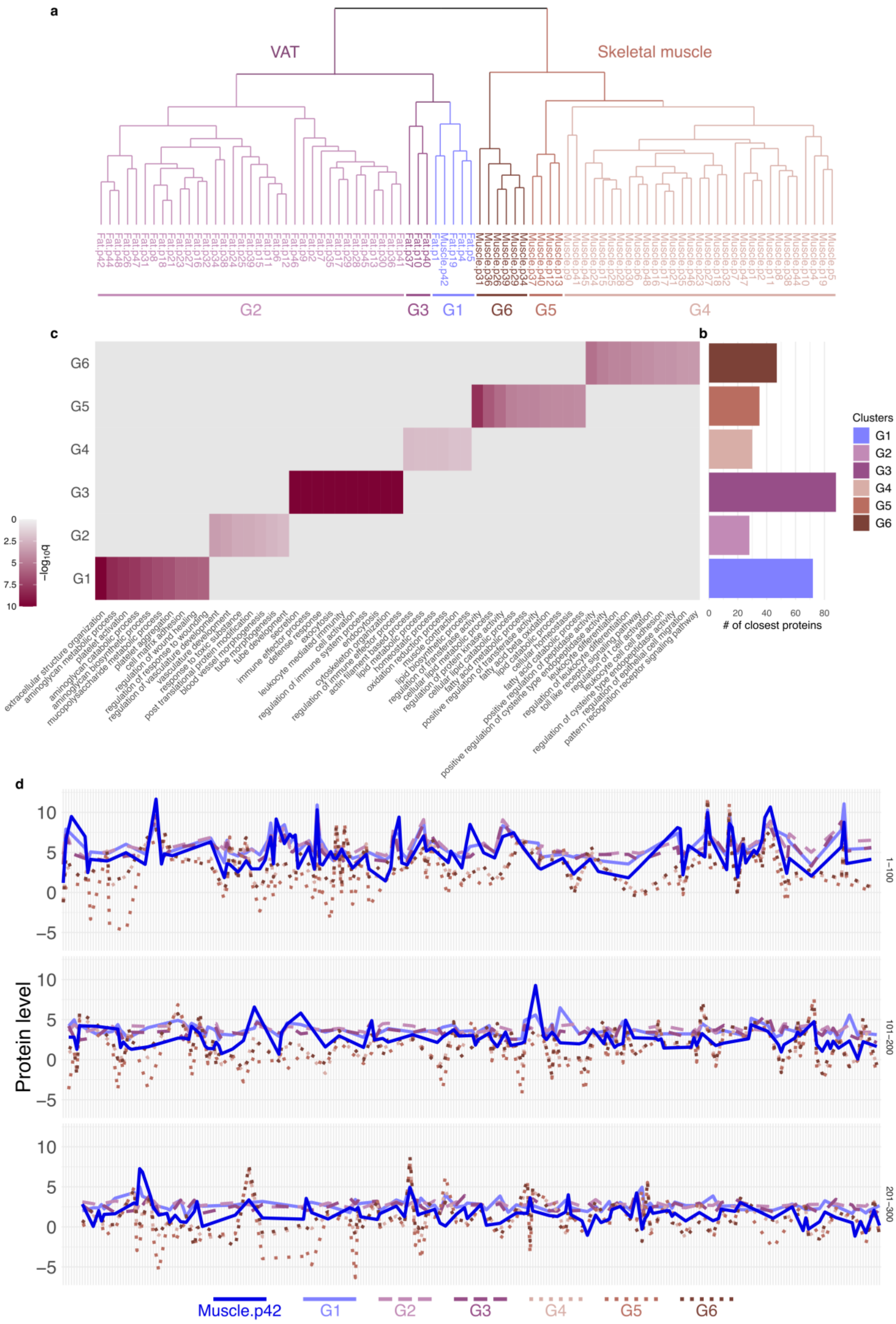
a
VAT
Skeletal muscle
G2
G3
G1
G6
G5
G4
c
b
G6
G5
G4
G3
G2
G1
−log10q
Clusters
G1
G2
G3
G4
G5
G6
### of closest proteins
d
Protein level
1–100
101–200
201–300
Muscle.p42
G1
G2
G3
G4
G5
G6

**Supplementary Figure S9: Investigation of the misclassified sample p42 from skeletal muscle. a)** Zoom into the hierarchical clustering (HC) from (Figure 1c) for VAT and skeletal muscle. **b)** Number of the top 300 most abundant proteins shared between VAT and skeletal muscle that are closer to sample p42 from skeletal muscle. Proteins for each group defined from HC have been collapsed to the average of their respective groups. **c)** Top 10 most significantly enriched GO terms from the GO enrichment analysis performed on the lists of proteins of each group in *b*. **d)** Expression of the top 300 most abundant proteins shared between VAT and skeletal muscle. p42 in skeletal muscle is shown independently while proteins from the other groups were collapsed to the average of the respective group.

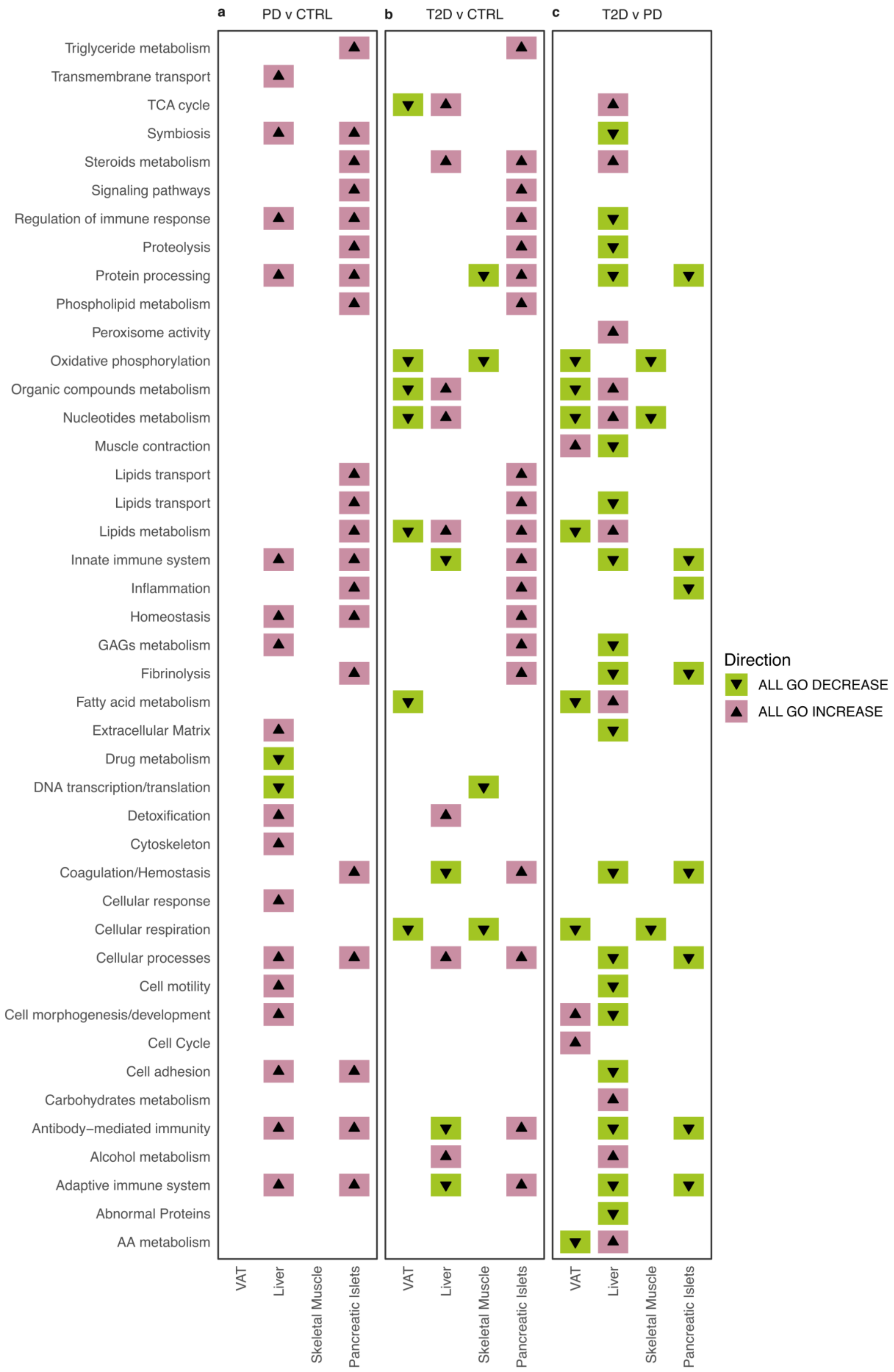

a
PD v CTRL
b
T2D v CTRL
c
T2D v PD
Triglyceride metabolism
Transmembrane transport
TCA cycle
Symbiosis
Steroids metabolism
Signaling pathways
Regulation of immune response
Proteolysis
Protein processing
Phospholipid metabolism
Peroxisome activity
Oxidative phosphorylation
Organic compounds metabolism
Nucleotides metabolism
Muscle contraction
Lipids transport
Lipids transport
Lipids metabolism
Innate immune system
Inflammation
Homeostasis
GAGs metabolism
Fibrinolysis
Fatty acid metabolism
Extracellular Matrix
Drug metabolism
DNA transcription/translation
Detoxification
Cytoskeleton
Coagulation/Hemostasis
Cellular response
Cellular respiration
Cellular processes
Cell motility
Cell morphogenesis/development
Cell Cycle
Cell adhesion
Carbohydrates metabolism
Antibody–mediated immunity
Alcohol metabolism
Adaptive immune system
Abnormal Proteins
AA metabolism
VAT
Liver
Skeletal Muscle
Pancreatic Islets
Direction
ALL GO DECREASE
ALL GO INCREASE

**Supplementary Figure S10: Direction of change for curated groups of GO terms for significantly enriched biological processes (q<0.01). a-c)** Panels represent pair-wise comparisons of phenotypes, columns represent tissues and rows represent curated groups of GO terms for biological processes. A combination of a triangle pointing upwards and a pink background color implies increase in the group, while those pointing downwards in combination with green background imply decrease. A graphic representation of the individual GO terms and as well as the curated groups is shown in (Figure 2), and detailed information on grouping and significance is shown in (Supplementary Table S8).

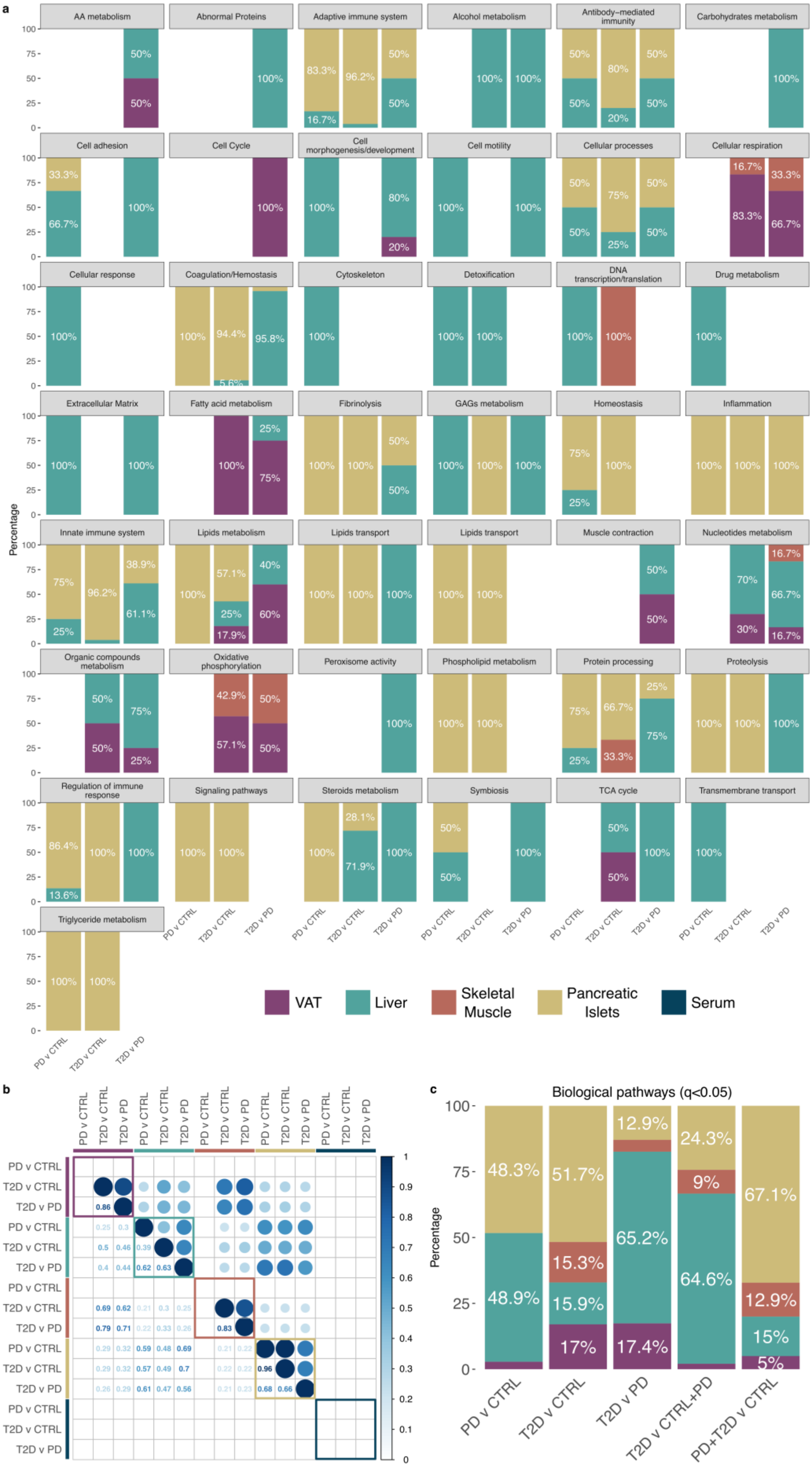
a
AA metabolism
Abnormal Proteins
Adaptive immune system
Alcohol metabolism
Antibody−mediated immunity
Carbohydrates metabolism
Cell adhesion
Cell Cycle
Cell morphogenesis/development
Cell motility
Cellular processes
Cellular respiration
Cellular response
Coagulation/Hemostasis
Cytoskeleton
Detoxification
DNA transcription/translation
Drug metabolism
Extracellular Matrix
Fatty acid metabolism
Fibrinolysis
GAGs metabolism
Homeostasis
Inflammation
Innate immune system
Lipids metabolism
Lipids transport
Lipids transport
Muscle contraction
Nucleotides metabolism
Organic compounds metabolism
Oxidative phosphorylation
Peroxisome activity
Phospholipid metabolism
Protein processing
Proteolysis
Regulation of immune response
Signaling pathways
Steroids metabolism
Symbiosis
TCA cycle
Transmembrane transport
Triglyceride metabolism
Percentage
PD v CTRL
T2D v CTRL
T2D v PD
VAT
Liver
Skeletal Muscle
Pancreatic Islets
Serum
b
PD v CTRL
T2D v CTRL
T2D v PD
c
Biological pathways (q<0.05)
Percentage
PD v CTRL
T2D v CTRL
T2D v PD
T2D v CTRL+PD
PD+T2D v CTRL

**Supplementary Figure S11: Distributions and similarities of enriched GO biological processes and biological pathways across tissues for pairwise comparisons of CTRL, PD, T2D, and their merged groups CTRL+PD and PD+T2D. a)** Distribution of enriched GO biological processes (q<0.01) across curated groupings and pair-wise phenotype comparisons among CTRL, PD and T2D. For groups of GO biological processes from (Figure 2) the fraction of terms from tissues of origin was calculated. Details on the groups are shown in (Supplementary Table S8). **b)** Similarity of GO biological terms for pair-wise comparisons of CTRL, PD and T2D across tissues**.** The similarity analysis was performed on sets of enriched GO biological processes (q<0.01) using the function *mgoSim* from the R package *GOSemSim* to obtain a combined similarity score with parameters *measure="Wang"* and *combine="BMA"*. **c)** Distribution of enriched biological pathways at 5% FDR corresponding to (Figure 3; Supplementary Table S9).

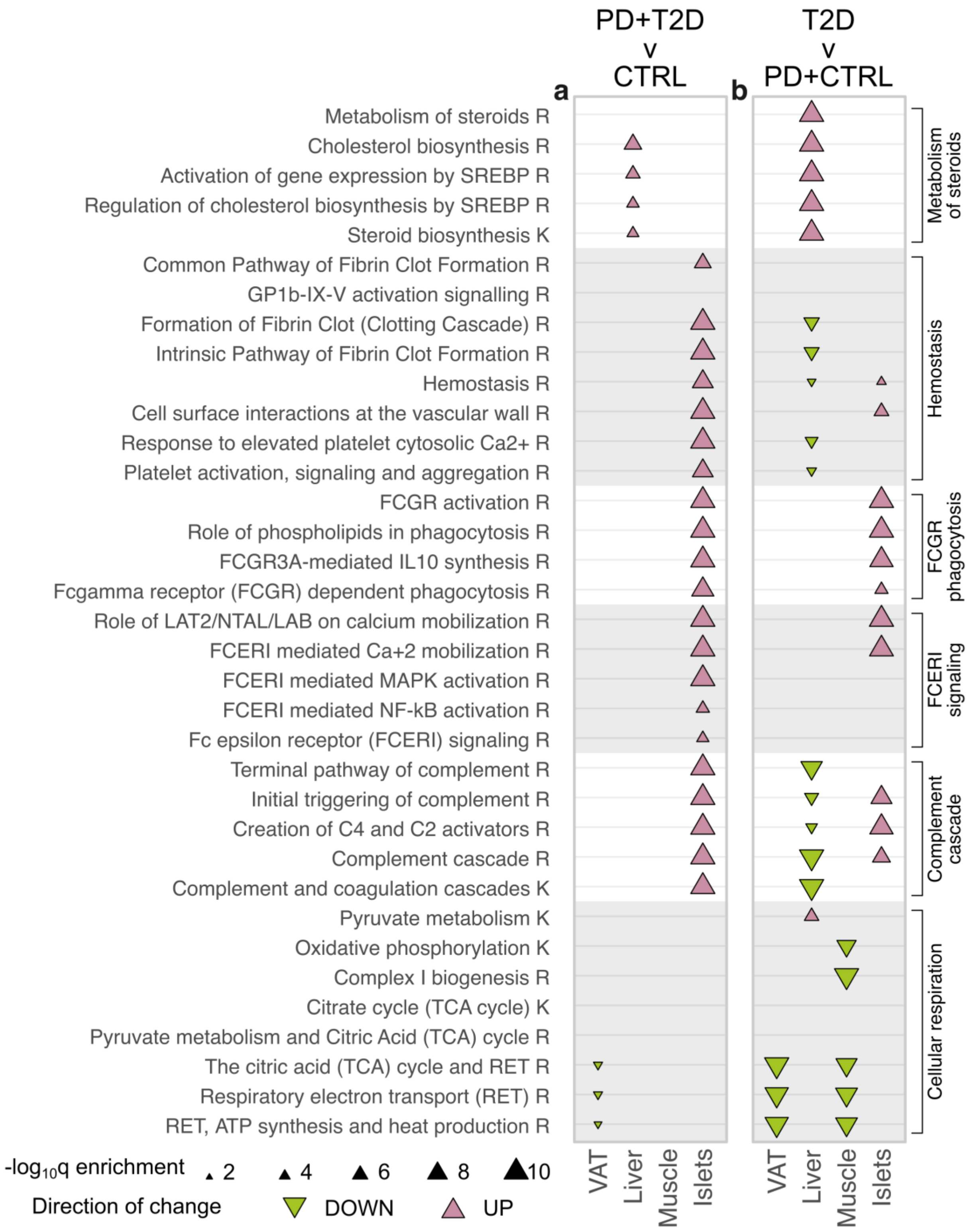


**Supplementary Figure S12: Selected groups of enriched biological pathways (q<0.05) for comparisons of the merged groups of CTRL+PD and PD+T2D to T2D and CTRL, respectively.** The plot contains an identical set to the one shown in (Figure 3). Rows represent pathways, panels represent pair-wise comparisons and columns within panels represent tissues. The database of origin for each pathway is noted with a suffix R or K implying Reactome or KEGG, respectively. A full list of enriched pathways is shown in (Supplementary Table S9). Sterol regulatory element binding proteins have been abbreviated as SREBP and Glycoprotein Ib-IX-V has been abbreviated as GP1b-IX-V.

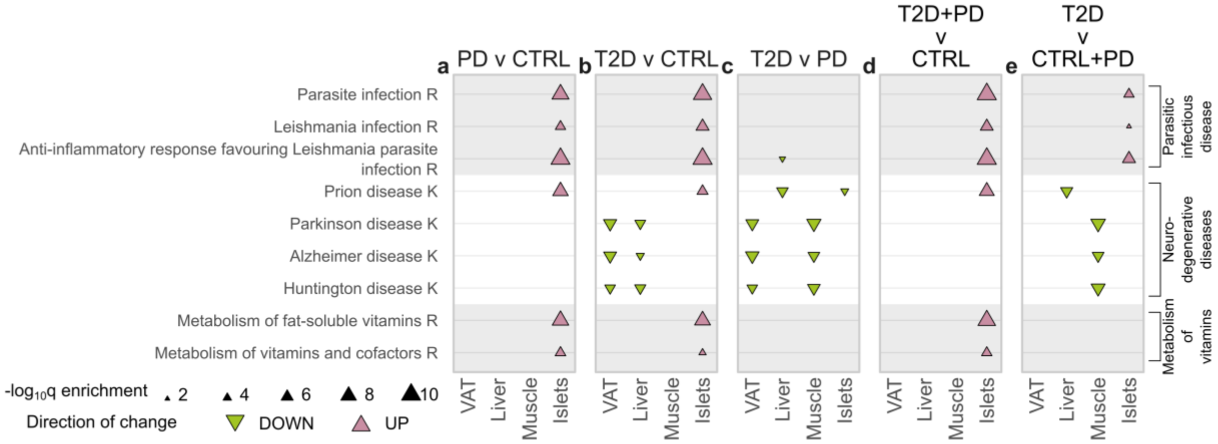


**Supplementary Table S13: Selected groups of enriched biological pathways (q<0.05) for pair-wise comparisons among CTRL, PD and T2D, and of the merged groups of CTRL+PD and PD+T2D to T2D and CTRL, respectively. a-e)** Rows represent pathways, panels represent pair-wise comparisons and columns within panels represent tissues. The database of origin for each pathway is noted with a suffix R or K implying Reactome or KEGG, respectively. A full list of enriched pathways is shown in (Supplementary Table S9).

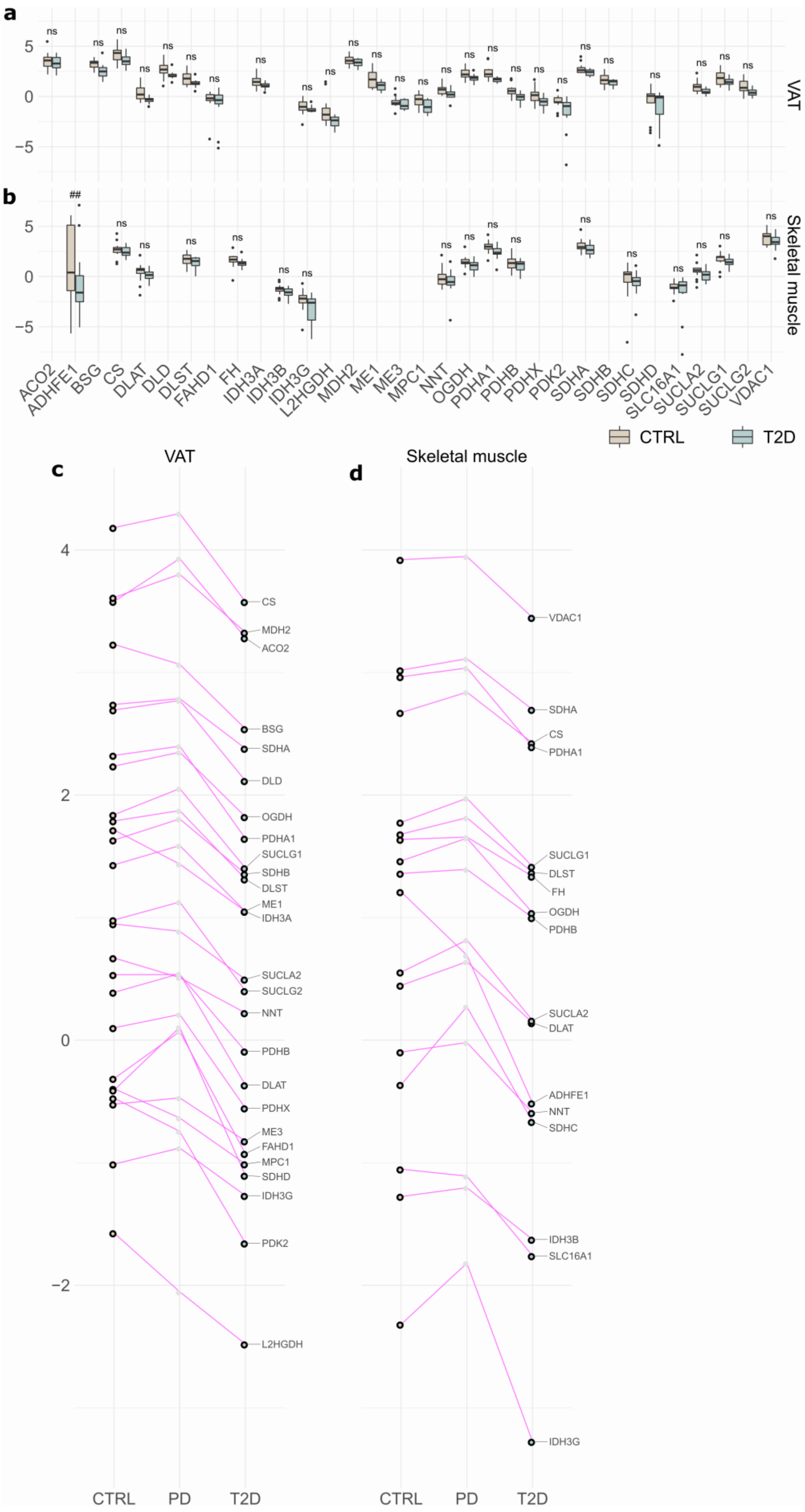
a
b
VAT
Skeletal muscle
ns
##
5
0
−5
ACO2
ADHFE1
BSG
CS
DLAT
DLD
DLST
FAHD1
FH
IDH3A
IDH3B
IDH3G
L2HGDH
MDH2
ME1
ME3
MPC1
NNT
OGDH
PDHA1
PDHB
PDHX
PDK2
SDHA
SDHB
SDHC
SDHD
SLC16A1
SUCLA2
SUCLG1
SUCLG2
VDAC1
CTRL
T2D
c
VAT
d
Skeletal muscle
4
2
0
−2
CS
MDH2
ACO2
BSG
SDHA
DLD
OGDH
PDHA1
SUCLG1
SDHB
DLST
ME1
IDH3A
SUCLA2
SUCLG2
NNT
PDHB
DLAT
PDHX
ME3
FAHD1
MPC1
SDHD
IDH3G
PDK2
L2HGDH
VDAC1
SDHA
CS
PDHA1
SUCLG1
DLST
FH
OGDH
PDHB
SUCLA2
DLAT
ADHFE1
NNT
SDHC
IDH3B
SLC16A1
IDH3G
CTRL
PD
T2D

**Supplementary Figure S14: Details on levels of proteins for TCA cycle and pyruvate metabolism in VAT and skeletal muscle for T2D and CTRL. a-b)** The two boxplots display differences in expression levels in **a)** VAT and **b)** skeletal muscle samples between CTRL and T2D. ***: $\pi \leq 2.1281$, **: $\pi \leq 1.1733$, *: $\pi \leq 0.6274$, #: $\pi \leq 0.4292$, ##: $\pi \leq 0.2572$, ns: $\pi > 0.2572$. **c-d)** The two bottom plots illustrate the overall trend of changes in expression of proteins characterized as driving for the TCA cycle and pyruvate metabolism pathway across CTRL, PD and T2D in **c)** VAT and **d)** skeletal muscle. Dots show average protein expression and a black circle around dots marks the outcomes that are being compared in the current example. Lines are used to mark alterations across phenotypes for a given protein.

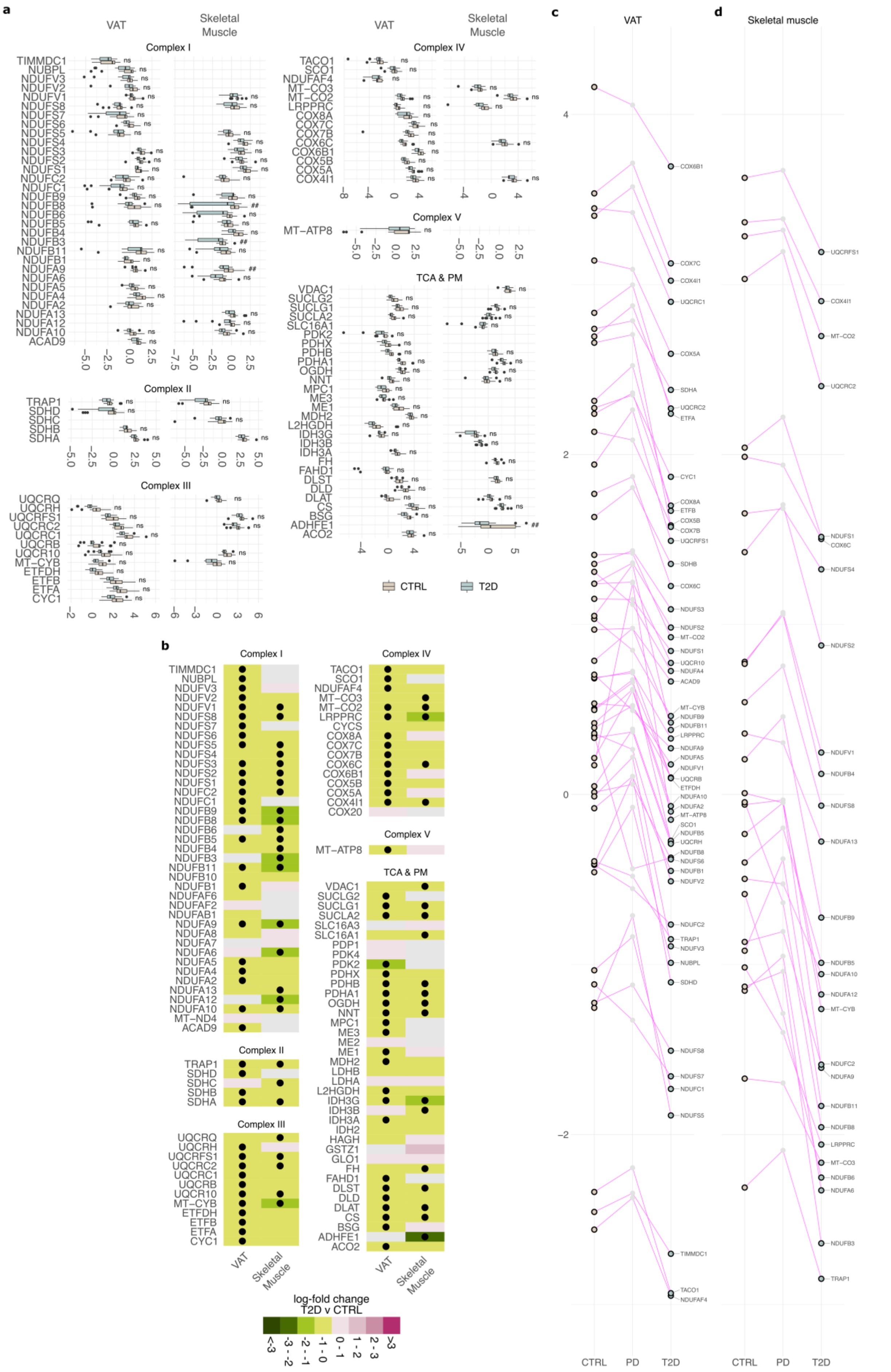

a
VAT
Skeletal Muscle
Complex I
Complex II
Complex III
Complex IV
Complex V
TCA & PM
CTRL
T2D
b
log-fold change T2D v CTRL
c
VAT
d
Skeletal muscle
CTRL
PD
T2D

**Supplementary Figure S15: Details on levels of proteins for respiratory electron transport, ATP synthesis by chemiosmotic coupling, and heat production by uncoupling proteins in VAT and skeletal muscle for T2D and CTRL. a-b)** Boxplot and heatmap were stratified based on mitochondrial protein complexes (Supplementary Table S11). **a)** Boxplots display differences in expression levels in VAT and skeletal muscle samples between CTRL and T2D. ***: $\pi \leq 2.1281$, **: $\pi \leq 1.1733$, *: $\pi \leq 0.6274$, #: $\pi \leq 0.4292$, ##: $\pi \leq 0.2572$, ns: $\pi > 0.2572$. **b)** Heatmap shows log-fold change of proteins identified in VAT and skeletal muscle. Black dots mark driving proteins in altering the pathway. Grey tiles mark non identified proteins in the respective tissue. **c-d)** Overall trend of changes in expression of proteins characterized as driving for the respiratory electron transport, ATP synthesis by chemiosmotic coupling, and heat production by uncoupling proteins pathway across CTRL, PD and T2D for **c)** VAT and **d)** skeletal muscle. Dots show average protein expression and a black circle around dots marks the outcomes that are being compared in the current example. Lines are used to mark alterations across phenotypes for a given protein.

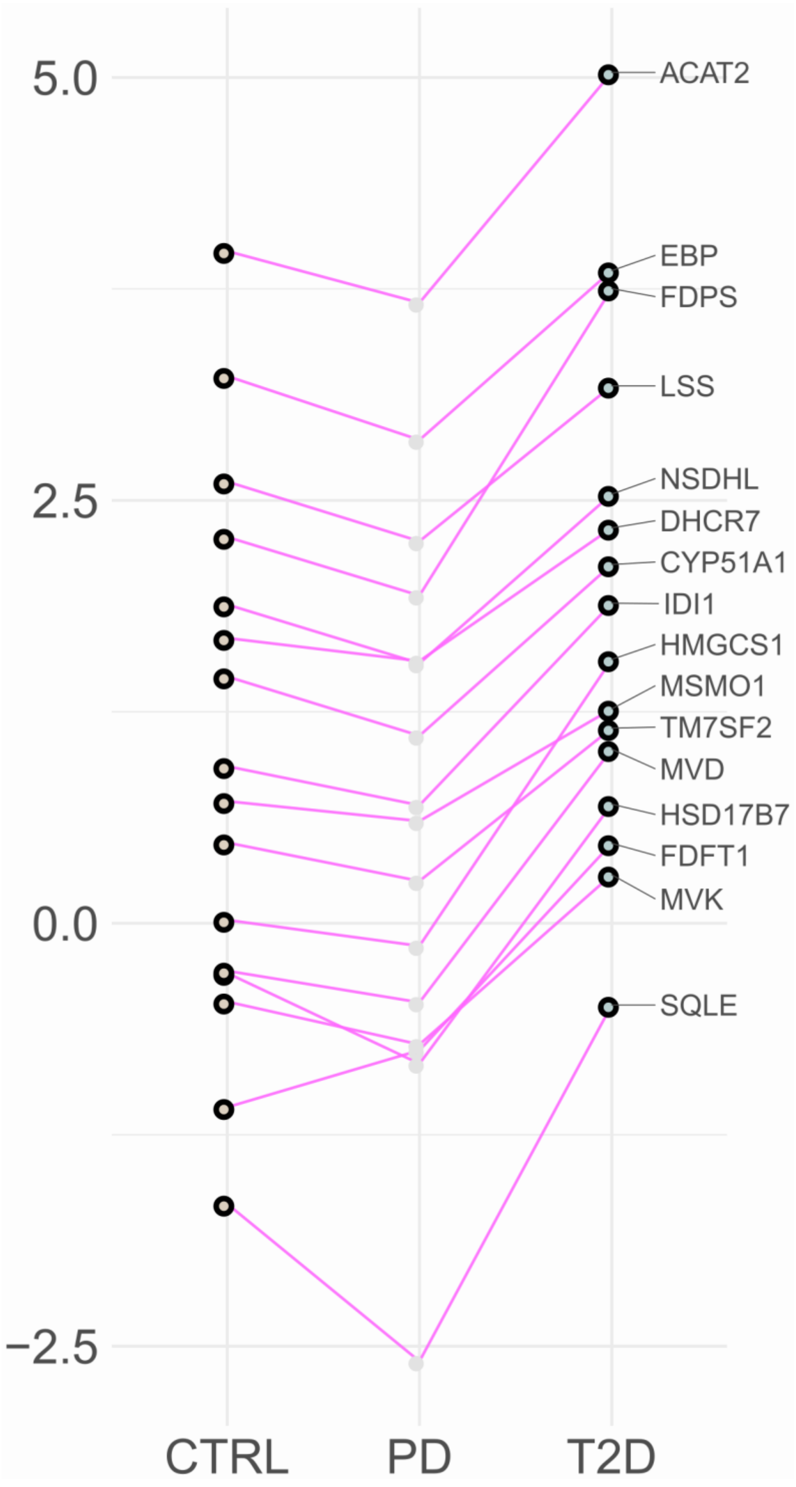


**Supplementary Figure S16: Overall trend of changes in expression of proteins characterized as driving for the cholesterol biosynthesis pathway in liver across CTRL, PD and T2D.** Dots show average protein expression and a black circle around dots marks the outcomes that are being compared in the current example. Lines are used to mark alterations across phenotypes for a given protein.

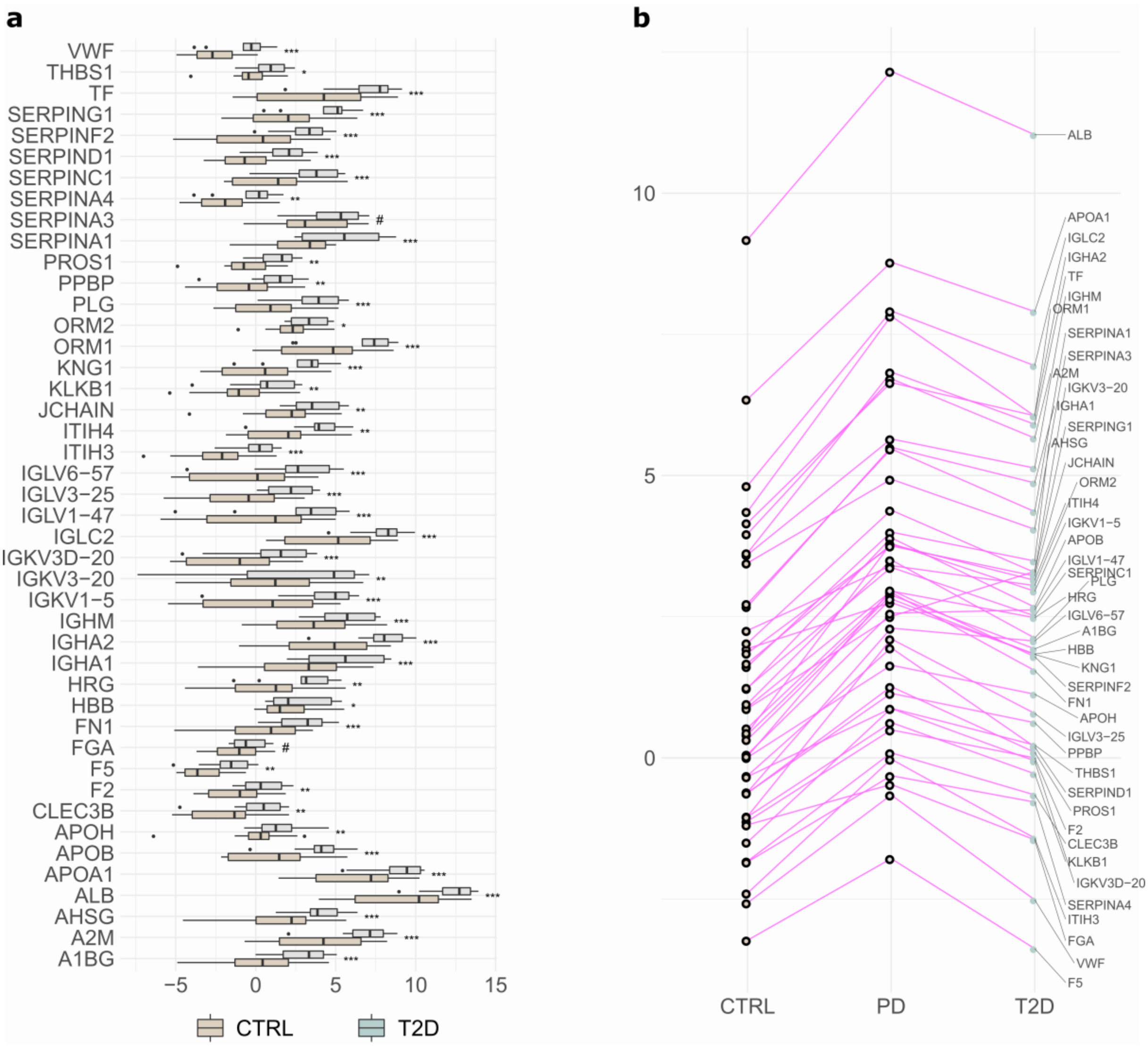


**Supplementary Figure S17: Details on levels of proteins for hemostasis in pancreatic islets for PD and CTRL. a)** Differences in expression levels between CTRL and PD. ***: $\pi \leq 2.1281$, **: $\pi \leq 1.1733$, *: $\pi \leq 0.6274$, #: $\pi \leq 0.4292$, ##: $\pi \leq 0.2572$, ns: $\pi > 0.2572$. **b)** Overall trend of changes in expression of proteins characterized as driving for the hemostasis pathway in pancreatic islets across CTRL, PD and T2D. Dots show average protein expression and a black circle around dots marks the outcomes that are being compared in the current example. Lines are used to mark alterations across phenotypes for a given protein.

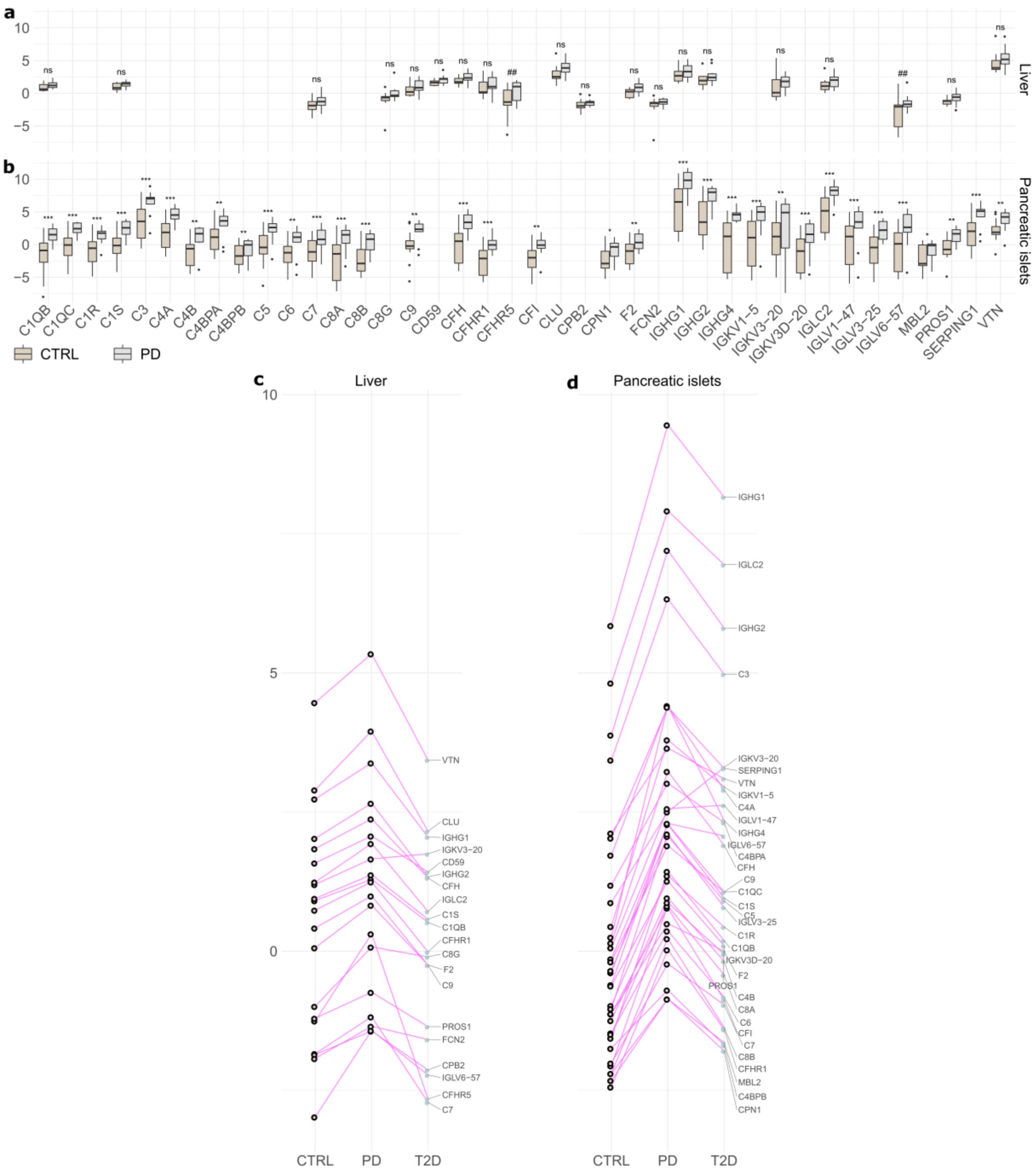


**Supplementary Figure S18: Details on levels of proteins for complement cascade in liver and pancreatic islets for PD and CTRL. a-b)** The boxplots display differences in expression levels from **a)** liver and **b)** pancreatic islets samples between CTRL and PD. ***: π≤2.1281, **: π≤1.1733, *: π≤0.6274, #: π≤0.4292, ##: π≤0.2572, ns: π>0.2572. **c-d)** Overall trend of changes in expression of proteins characterized as driving for the complement cascade pathway across CTRL, PD and T2D in **c)** liver and **d)** pancreatic islets. Dots show average protein expression and a black circle around dots marks the outcomes that are being compared in the current example. Lines are used to mark alterations across phenotypes for a given protein.
